# In-depth characterization of denitrifier communities across different soil ecosystems in the tundra

**DOI:** 10.1101/2020.12.21.419267

**Authors:** Igor S. Pessi, Sirja Viitamäki, Anna-Maria Virkkala, Eeva Eronen-Rasimus, Tom O. Delmont, Maija E. Marushchak, Miska Luoto, Jenni Hultman

**Affiliations:** Department of Microbiology, University of Helsinki, Helsinki, Finland; Helsinki Institute of Sustainability Science (HELSUS), Helsinki, Finland; Department of Geosciences and Geography, University of Helsinki, Helsinki, Finland; Woodwell Climate Research Center, Falmouth, MA, USA; Marine Research Centre, Finnish Environment Institute (SYKE), Helsinki, Finland; Department of Bioinformatics, Genoscope, Paris, France; Department of Biological and Environmental Science, University of Jyväskylä, Jyväskylä, Finland; Department of Environmental and Biological Sciences, University of Eastern Finland, Kuopio, Finland; Natural Resources Institute Finland (Luke), Helsinki, Finland

**Keywords:** Arctic, denitrification, genome-resolved metagenomics, nitrous oxide

## Abstract

**Background:** In contrast to earlier assumptions, there is now mounting evidence for the role of tundra soils as important sources of the greenhouse gas nitrous oxide (N_2_O). However, the microorganisms involved in the cycling of N_2_O in this system remain largely uncharacterized. Since tundra soils are variable sources and sinks of N_2_O, we aimed at investigating differences in community structure across different soil ecosystems in the tundra.

**Results:** We analysed 1.4 Tb of metagenomic data from soils in northern Finland covering a range of ecosystems from dry upland soils to water-logged fens and obtained 796 manually binned and curated metagenome-assembled genomes (MAGs). We then searched for MAGs harbouring genes involved in denitrification, an important process driving N_2_O emissions. Communities of potential denitrifiers were dominated by microorganisms with truncated denitrification pathways (i.e., lacking one or more denitrification genes) and differed across soil ecosystems. Upland soils showed a strong N_2_O sink potential and were dominated by members of the Alphaproteobacteria such as *Bradyrhizobium* and *Reyranella*. Fens, which had in general net-zero N_2_O fluxes, had a high abundance of poorly characterized taxa affiliated with the Chloroflexota lineage Ellin6529 and the Acidobacteriota subdivision Gp23.

**Conclusions:** By coupling an in-depth characterization of microbial communities with *in situ* measurements of N_2_O fluxes, our results suggest that the observed spatial patterns of N_2_O fluxes in the tundra are related to differences in the composition of denitrifier communities.

## Background

Nitrous oxide (N_2_O) is a greenhouse gas (GHG) that has approximately 300 times the global warming potential of carbon dioxide on a 100-year scale [1]. Atmospheric N_2_O concentrations have increased by nearly 20% since pre-industrial times, with soils – both natural and anthropogenic – accounting for up to 70% of the global emissions [2]. Despite being nitrogen (N) limited and enduring low temperatures throughout most of the year, tundra soils are increasingly recognized as important sources of N_2_O [3–7]. The relative contribution of tundra soils to global GHG emissions is predicted to increase in the future [8, 9], as the warming rate at high latitude environments is more than twice as high than in other regions [10].

Microbial denitrification is an important source of N_2_O [11]. Denitrification is a series of enzymatic steps in which nitrate (NO_3_^-^) is sequentially reduced to nitrite (NO_2_-), nitric oxide (NO), N_2_O, and dinitrogen (N2) via the activity of the Nar, Nir, Nor, and Nos enzymes, respectively. The denitrification trait is common across a wide range of archaea, bacteria, and some fungi, most of which are facultative anaerobes that switch to N oxides as electron acceptor when oxygen becomes limiting [12]. Denitrification is a modular community process performed in synergy by different microbial taxa that execute only a subset of the complete denitrification pathway [12, 13]. With the growing number of microbial genomes sequenced in recent years, it has become evident that only a fraction of the microorganisms involved in the denitrification pathway encode the enzymatic machinery needed for complete denitrification [14, 15].

Compared to high N_2_O-emitting systems such as agricultural and tropical soils, our knowledge of denitrifier communities in tundra soils is limited. As denitrification leads to the loss of N to the atmosphere, it enhances the N-limited status of tundra systems thus impacting both microbial and plant communities [16, 17]. Investigations of denitrifier diversity in the tundra have been largely limited to gene-centric surveys using microarrays, amplicon sequencing, qPCR, and read-based metagenomics, which provide limited information on the taxonomic identity and genomic composition of community members. These studies have shown that denitrifier communities in the tundra are dominated by members of the phyla Proteobacteria, Actinobacteria, and Bacteroidetes, and that the potential for complete denitrification is usually present at the community level [18–22]. However, it is not known whether the complete denitrification potential occurs within discrete microbial populations or is widespread throughout populations of truncated denitrifiers lacking one or more denitrification genes. In addition, tundra soils encompass many different ecosystems, some of which are notorious N_2_O sources (e.g. bare peat surfaces [3]). N_2_O consumption is usually favoured in wetlands, where low NO_3_^-^ availability due to anoxia promotes the reduction of N_2_O to N_2_ [23]. In upland soils, N_2_O fluxes vary in both time and space. Strong N_2_O sinks have been observed specially in sparsely vegetated upland soils [7], but the microbial processes underlying N_2_O consumption in these systems are largely unknown [24]. Altogether, these large differences in N_2_O fluxes across tundra ecosystems indicate differences in the structure of microbial communities, but a comprehensive understanding of the microorganisms driving N_2_O fluxes in tundra soils is lacking.

Modelling N_2_O emissions based on microbial community structure is challenging. N_2_O fluxes are characterized by a high temporal and spatial heterogeneity driven by several environmental constraints related to soil pH, N, moisture, and oxygen content [11]. In addition, our knowledge of the regulation of the denitrification process is largely based on the activity of model organisms such as the complete denitrifier *Paracoccus denitrificans* [25]. It has been suggested that incomplete denitrifiers that contain Nir and Nor but lack Nos contribute substantially to soil N_2_O emissions [26], while non-denitrifying N_2_O reducers, i.e. microorganisms that contain Nos but lack Nir, can represent an important N_2_O sink [27–29]. Furthermore, the partitioning of metabolic pathways across different populations with truncated pathways – also known as metabolic handoffs [30] – has been linked to higher efficiencies in substrate consumption compared to complete pathways [15, 31]. However, it remains largely unclear how populations of truncated denitrifiers with different sets of denitrification genes interact with each other and the environment impacting *in situ* N_2_O emissions.

The paucity of in-depth knowledge on denitrifying communities in the tundra impairs our ability to model current and future N_2_O fluxes from this biome. A better understanding of the ecological, metabolic, and functional traits of denitrifiers is thus critical for improving current models and mitigating N_2_O emissions [32]. This invariably relies on the characterization of the so-called uncultured majority, i.e. microorganisms that have not been cultured to date but which comprise a high proportion of the microbial diversity in complex ecosystems [33, 34]. Genome-resolved metagenomics is a powerful tool to access the genomes of uncultured microorganisms and has provided important insights into carbon cycling processes in tundra soils [35–37]. However, this approach has not yet been applied to investigate the mechanisms driving N_2_O fluxes in the tundra. Here, we used genome-resolved metagenomics to investigate the diversity and metabolic capabilities of denitrifiers across different tundra soil ecosystems characterised by a high variability in net N_2_O fluxes in an area of mountain tundra in Kilpisjärvi, northern Finland.

## Methods

### Study area and sampling

The Saana Nature Reserve (69.04°N, 20.79°E) is located in Kilpisjärvi, northern Finland **(Fig. 1a)**. The area is part of the mountain tundra biome and is characterized by a mean annual temperature of –1.9°C and annual precipitation of 487 mm [38]. Our study sites are distributed across Mount Saana and Mount Korkea-Jehkas and the valley in between **(Fig. 1b)**, and include barren soils, heathlands (dominated by evergreen and deciduous shrubs), meadows (dominated by graminoids and forbs), and fens **(Fig. 1c)**. Sampling was performed across 43 sites (barren soils, n = 2; heathlands, n = 18; meadows, n = 7; fens, n = 16) during the peak of the growing season in the northern hemisphere. Fen sites were sampled in July 2018 and all other sites in July 2017. Samples were obtained with a soil corer sterilized with 70% ethanol and, when possible, cores were split into organic and mineral samples using a sterilized spatula. In total, 69 samples (41 organic and 28 mineral) were obtained from the 43 sites **(Figure 1c, Suppl. Table S1)**. Samples were transferred to a whirl-pack bag and immediately frozen in dry ice. Samples were transported frozen to the laboratory at the University of Helsinki and kept at –80°C until analyses.

**Fig. 1.**
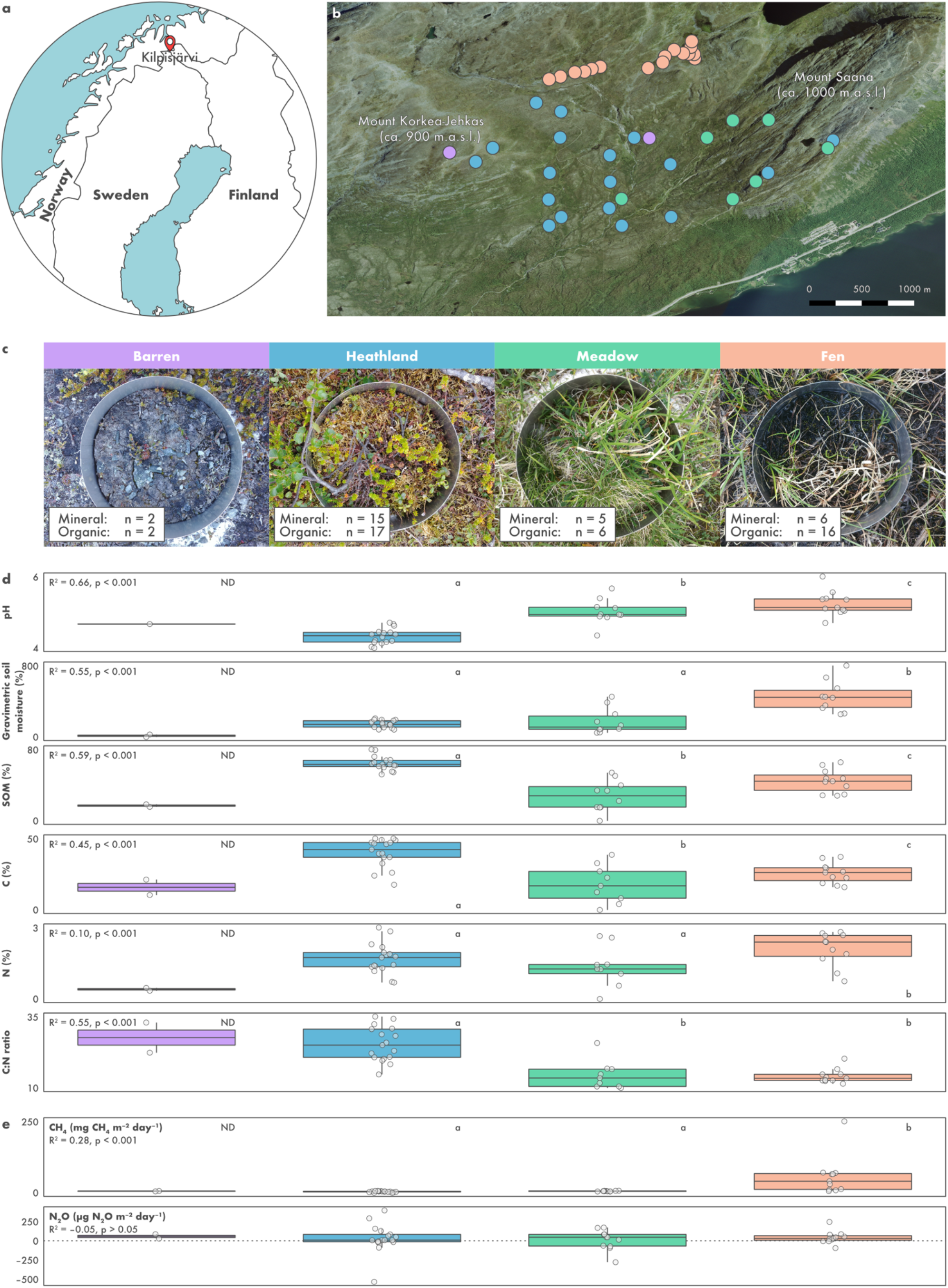
Saana Nature Reserve, an area of mountain tundra in Kilpisjärvi, northern Finland. **a)** Map of Fennoscandia showing the location of Kilpisjärvi. **b)** Aerial overview of the study area showing the location of the 43 sites sampled for metagenomic analysis. Image provided by the National Land Survey of Finland under the Creative Commons CC BY 4.0 license. **c)** *In situ* photographs of the four types of soil ecosystems investigated. **d)** Physicochemical characterization of the soil ecosystems based on 41 organic and 28 mineral samples taken from the 43 sites. **e)** *In situ* ecosystem-level nitrous oxide (N_2_O) and methane (CH_4_) fluxes measured from the 43 sites using a static, non-steady state, non-flow-through system. Negative values represent net uptake and positive net emissions. For clarity, one outlier measurement from a meadow site (660 μg N_2_O m^-2^ day^-1^) was removed. In panels d and e, ecosystems followed by different letters are significantly different (one-way ANOVA, p < 0.05). Samples from barren soils were not included in the ANOVA procedure due to the limited number of samples (ND: not determined).

### Soil physicochemical characterization and *in situ* measurement of GHG fluxes

Soil pH, moisture, and soil organic matter (SOM) content were measured from the 69 samples according to Finnish (SFS) and international (ISO) standards (SFS 300, ISO 10390, and SFS 3008). Carbon (C) and N content were measured using a Vario Micro Cube machine (Elementar, Langenselbold, Germany). *In situ* ecosystem-level N_2_O and methane (CH_4_) fluxes were measured from the 43 sites using a static, non-steady state, non-flow-through system composed of a darkened acrylic chamber (20 cm diameter, 25 cm height) [4, 39]. Measurements were conducted between 2^nd^ July and 2^nd^ August 2018, between 10 am and 5 pm. Simultaneous measurement of GHG fluxes and sampling for metagenomic sequencing was not possible due to limited resources and logistic constraints. At each site, five 25 mL gas samples were taken during a 50-minute chamber closure and transferred to evacuated Exetainer vials (Labco, Lampeter, UK). Gas samples were analysed using an Agilent 7890B gas chromatograph (Agilent Technologies, Santa Clara, CA, USA) equipped with an autosampler (Gilson, Middleton, WI, USA) and a flame ionization detector for CH_4_ and an electron capture detector for N_2_O. Gas concentrations were calculated from the gas chromatograph peak areas based on standard curves with a CH_4_ concentration of 0–100 ppm and a N_2_O concentration of 0–5000 ppb. Differences in physicochemical composition and rates of GHG fluxes across soil ecosystems were assessed using one-way analysis of variance (ANOVA) followed by Tukey’s HSD test with the *lm* and *TukeyHSD* functions in R v3.6.3 (https://www.r-project.org). Due to the limited number of samples from barren sites, these were not included in the ANOVA procedure.

### Metagenome sequencing and processing of raw data

Total DNA and RNA were co-extracted as previously described [40]. Briefly, extraction was performed on 0.5 g of soil using a hexadecyltrimethyl ammonium bromide (CTAB), phenol-chloroform, and bead-beating protocol. DNA was purified using the AllPrep DNA Mini Kit (QIAGEN, Hilden, Germany) and quantified using the Qubit dsDNA BR Assay Kit (ThermoFisher Scientific, Waltham, MA, USA). Library preparation for Illumina metagenome sequencing was performed using the Nextera XT DNA Library Preparation Kit (Illumina, San Diego, CA, USA). Metagenomes were obtained for 69 samples across two paired-end NextSeq (132–170 bp) and one NovaSeq (2 x 151 bp) runs **(Suppl. Table S1)**. Two samples were additionally sequenced with Nanopore MinION. For this, libraries were prepared using the SQK-LSK109 Ligation Sequencing Kit with the long fragment buffer (Oxford Nanopore Technologies, Oxford, UK) and the NEBNext Companion Module for Oxford Nanopore Technologies Ligation Sequencing Kit (New England Biolabs). Each sample was sequenced for 48 hours on one R9.4 flow cell.

We obtained more than 9 billion Illumina (1.4 Tb) and 7 million Nanopore (21.5 Gb) reads from the 69 soil metagenomes (mean: 19.9 Gb, minimum: 0.7 Gb, maximum: 82.9 Gb) **(Suppl. Table S1)**. The quality of the raw Illumina data was verified with fastQC v0.11.9 (https://www.bioinformatics.babraham.ac.uk/projects/fastqc) and multiQC v1.8 [41]. Cutadapt v1.16 [42] was then used to trim sequencing adapters and low-quality base calls (q < 20) and to filter out short reads (< 50 bp). Nanopore data were basecalled with GPU guppy v4.0.11 using the high-accuracy model and applying a minimum quality score of 7. The quality of the basecalled Nanopore data was assessed with pycoQC v2.5.0.21 [43] and adapters were trimmed with Porechop v0.2.4 (https://github.com/rrwick/Porechop).

### Taxonomic profiling

Taxonomic profiles of the microbial communities were obtained using a read-based approach, i.e., based on unassembled Illumina data. Due to differences in sequencing depth across the samples, the dataset was resampled to 2,000,000 reads per sample with seqtk v1.3 (https://github.com/lh3/seqtk). Reads matching the SSU rRNA gene were identified with METAXA v2.2 [44] and classified against the SILVA database release 138.1 [45] in mothur v1.44.3 [46] using the Wang’s Naïve Bayesian Classifier [47] and a 80% confidence cut-off. Differences in community structure were assessed using non-metric multidimensional scaling (NMDS) and permutational ANOVA (PERMANOVA) with the package vegan v2.5.6 (https://cran.r-project.org/web/packages/vegan) in R v3.6.3 (functions *metaMDS* and *adonis*, respectively). Relationships between the abundance of individual genera and N_2_O flux rates were assessed using linear regression in R v3.6.3 (https://www.r-project.org).

### Metagenome assembling and binning

Metagenome assembling of the Illumina data was performed as two co-assemblies. One co-assembly comprised the upland soils (barren, heathland, and meadow; n = 47) and the other the fen samples (n = 22). For each co-assembly, reads from the respective samples were pooled and assembled with MEGAHIT v1.1.1.2 [48]. Assembling of the Nanopore data was done for each sample individually with metaFlye v2.7.1 [49], and contigs were corrected based on Illumina data from the respective sample with bowtie v2.3.5 [50], SAMtools v1.9 [51], and pilon v1.23 [52]. Quality assessment of the (co-)assemblies was obtained with metaQUAST v5.0.2 [53].

MAG binning was done separately for each Illumina and Nanopore (co-)assembly with anvi’o v6.2 [54] after discarding contigs shorter than 2500 bp. The two Illumina co-assemblies and the two individual Nanopore assemblies yielded more than 4 million contigs longer than 2,500 bp, with a total assembly size of 21.1 Gb. Gene calls were predicted with prodigal v2.6.3 [55]. Single-copy genes were identified with HMMER v.3.2.1 [56] and classified with DIAMOND v0.9.14 [57] against the Genome Taxonomy Database (GTDB) release 04-RS89 [58, 59]. Illumina reads were mapped to the contigs with bowtie v2.3.5 [50] and SAM files were sorted and indexed using SAMtools v1.9 [51]. The co-assemblies covered a significant fraction of the original metagenomic data, with an average read recruitment rate of 54.6% across samples (minimum: 22.9%, maximum: 75.8%). Due to their large sizes, Illumina co-assemblies were split into 100 smaller clusters based on differential coverage and tetranucleotide frequency with CONCOCT v1.0.0 [60]. Contigs were then manually sorted into bins based on the same composition and coverage metrics using the *anvi-interactive* interface in anvi’o v6.2 [54]. Nanopore contigs were binned directly without pre-clustering. Bins that were ≥ 50% complete according to the presence of single-copy genes were further refined using the *anvi-refine* interface in anvi’o v6.2 [54]. In addition to taxonomic signal (based on single-copy genes classified against GTDB), either differential coverage or tetranucleotide frequency was used to identify and remove outlying contigs. The former was used for bins with a large variation in contig coverage across samples, and the latter for those with marked differences in GC content across contigs. Medium- and high-quality bins (≥ 50% complete and < 10% redundant according to the MIMAG standard [61]) were renamed as MAGs and kept for downstream analyses.

### Gene-centric analyses

Functional profiles of the microbial communities were obtained using a gene-centric approach based on assembled data. For each (co-)assembly, gene calls were translated to amino acid sequences and searched against the KOfam hidden Markov model (HMM) database with KofamScan v1.3.0 [62]. Only matches with scores above the pre-computed family-specific thresholds were kept. Genes putatively identified as denitrification genes (*nirK, nirS, norB*, and *nosZ*) were submitted to further analyses to identify false positives consisting of distant homologues that are not involved in denitrification. Amino acid sequences were aligned with MAFFT v7.429 [63] and alignments were visualized with Unipro UGENE v38.1 [64]. Sequences were then inspected for the presence of conserved residues at positions associated with the binding of co-factors and active sites: *nirK*, Cu-binding and active sites [65]; *nirS*, c-heme and d_1_-heme binding sites [66]; *norB*, binding of the catalytic centres cyt b, b_3_, and Feb [67]; *nosZ*: binding of the Cu_Z_ and Cu_A_ centres [67]. Sequences which did not contain the correct amino acid at these positions were removed. Finally, resulting amino acid sequences were aligned with MAFFT v7.429 [63] along with reference sequences from the genome of cultured denitrifiers [14] and a maximum-likelihood tree was computed with FastTree v2.1.11 [68] using the LG+GAMMA model. Annotation of denitrification genes was also performed for previously published genomes retrieved from GenBank. These included a set of 1529 MAGs obtained from soils in Stordalen Mire, northern Sweden [37], and all (n = 69) genomes of Acidobacteriota strains and candidate taxa (accessed on 9 October 2020).

The abundance of functional genes was computed based on read coverage with CoverM v0.6.1 [69]. For this, Illumina reads were mapped to the contigs with minimap v2.17 [70] and coverage was normalized to reads per kilobase million (RPKM). Differences in functional community structure were assessed using NMDS and PERMANOVA as described above for the taxonomic profiles. Differences in the abundance of individual genes across soil ecosystems were assessed using ANOVA followed by Tukey’s HSD test with the *lm* and *TukeyHSD* functions in R v3.6.3 (https://www.r-project.org). Due to the limited number of samples from barren sites, these were not included in the ANOVA procedure. Relationships between the abundance of denitrification genes and N_2_O flux rates were assessed using linear regression in R v3.6.3.

### Phylogenomic analyses of MAGs and metabolic reconstruction

Phylogenetic placement of MAGs was done based on 122 archaeal and 120 bacterial single-copy genes with GTDB-Tk v1.3.0 [71] and the GTDB release 05-RS95 [58, 59]. Acidobacteriota MAGs containing denitrification genes were submitted to further phylogenomic analyses alongside all genomes of Acidobacteriota strains and candidate taxa available on GenBank (n = 69; accessed on 9 October 2020). For this, the amino acid sequence of 23 ribosomal proteins was retrieved for each genome with anvi’o v6.2 [54] and aligned with MUSCLE v3.8.1551 [72]. A maximum likelihood tree was then computed based on the concatenated alignments with FastTree v2.1.11 using the LG+GAMMA model [68]. *Escherichia coli* ATCC 11775 was used to root the tree.

For metabolic reconstruction, MAGs were annotated against the KOfam HMM database [62] with HMMER v.3.2.1 [56] using the pre-computed score thresholds of each HMM profile. The *anvi-estimate-metabolism* program in anvi’o v6.2 [54] was then used to predict the metabolic capabilities of the MAGs. A metabolic pathway was considered present in MAGs containing at least 75% of the genes involved in the pathway. Carbohydrate-active enzymes (CAZymes) were annotated with dbCAN v.2.0 based on the dbCAN v7 HMM database [73]. Only hits with an e-value < 1 x 10^-14^ and coverage > 0.35 were considered.

### MAG dereplication and read recruitment analysis

Prior to read recruitment analyses, Illumina and Nanopore MAGs were dereplicated based on a 99% average nucleotide identity (ANI) threshold with fastANI v1.3 [74] to remove redundancy (i.e., MAGs that were recovered multiple times across the different assemblies). Read recruitment analyses were then performed with CoverM v0.6.1 (https://github.com/wwood/CoverM). For this, Illumina reads were mapped to the set of non-redundant MAGs with minimap v2.17 [70] and relative abundances were calculated as a proportion of the reads mapping to each MAG.

## Results

### Environmental characterization and *in situ* GHG fluxes

Our sampling design in Kilpisjärvi included two soil depths across four ecosystems that are characteristic of the tundra biome (barren soils, heathlands, meadows, and fens) **(Fig. 1a–c)**. In previous studies, we have established in the area a systematic fine-scale sampling of microclimate, soil conditions, and vegetation in topographically distinct environments [40, 75, 76]. Local variation in topography and soil properties creates a mosaic of habitats characterized by contrasting ecological conditions. This makes the study setting ideal to investigate species-environment relationships and ecosystem functioning in the tundra [40, 77, 78]. Soil ecosystems differed in vegetation cover and physicochemical composition, with fens being characterized by higher pH, moisture, and N content (one-way ANOVA, R^2^ = 0.10–0.66, p < 0.001) and, together with the meadows, lower C:N ratio (one-way ANOVA, R^2^ = 0.55, p < 0.001) **(Fig. 1d)**.

*In situ* measurements showed a high sink-source variability in net N_2_O fluxes across the ecosystems **(Fig. 1e)**. Although the average N_2_O flux across all sites was small (net consumption of 6 μg N_2_O m^-2^ day^-1^), high N_2_O emission at rates of up to 660 μg N_2_O m^-2^ day^-1^ was observed at the meadow sites. Likewise, strong N_2_O consumption (up to –435 μg N_2_O m^-2^ day^-1^) was observed particularly at the heathland and meadow sites. Net CH_4_ emissions were observed exclusively at the fen sites **(Fig. 1e)**.

### Differences in microbial community structure across soils ecosystems

Read-based analyses of unassembled SSU rRNA gene sequences showed that microbial community composition differed across the ecosystems, with fen soils harbouring contrasting microbial communities compared to the other ecosystems (PERMANOVA, R^2^ = 0.35, p < 0.001) **(Suppl. Fig. S1a)**. No differences in community structure were observed between soil depths or the interaction between soil ecosystem and depth (PERMANOVA, p > 0.05). Among previously described (i.e., not unclassified) taxa, microbial communities in barren, heathland, and meadow soils were dominated by aerobic and facultative anaerobic heterotrophs such as *Acidipila/Silvibacterium, Bryobacter, Granulicella, Acidothermus, Conexibacter, Mycobacterium, Mucilaginibacter, Bradyrhizobium*, and *Roseiarcus* **(Suppl. Fig. S1b)**. On the other hand, fen soils were dominated by methanogenic archaea from the genera *Methanobacterium* and *Methanosaeta* and anaerobic bacteria such as *Thermoanaerobaculum*, *Desulfobacca*, and *Smithella*, but also the putative aerobic heterotroph *Candidatus* Koribacter. We did not observe a significant relationship between the abundance of individual microbial genera and N_2_O flux rates (linear regression, p > 0.05).

Communities from different ecosystems also differed in their functional potential **(Suppl. Fig. S1c).** Denitrification genes (*nirK*, *nirS*, *norB*, and *nosZ*) were in general more abundant in the meadows and fens (one-way ANOVA, R^2^ = 0.48–0.76, p < 0.001) **(Suppl. Fig. S1d)**. Fen soils, which had the highest gravimetric soil water content, also had a higher abundance of genes involved in sulfate reduction (*dsrA* and *dsrB*) and methanogenesis (*mcrA* and *mcrB*) (one-way ANOVA, R^2^ = 0.59–0.90, p < 0.001), indicating the prevalence of anoxic and reductive soil conditions in these wet sites. We did not observe a significant relationship between N_2_O flux rates and neither the abundance of individual denitrification genes nor the ratio between *nosZ* and *nirK*+*nirS* abundances (linear regression, p > 0.05). However, the ratio between *nosZ* and *nirK+nirS* abundances was higher in the meadows (one-way ANOVA, R^2^ = 0.29, p < 0.001) **(Suppl. Fig. S1d)**, which indicates a higher potential for N_2_O consumption in this ecosystem.

### A manually curated genomic database from tundra soil metagenomes

Using anvi’o [54], we obtained 8,043 genomic bins and manually curated these to a set of 796 medium- and high-quality MAGs (≥ 50% complete and ≤ 10% redundant according to the MIMAG standard [61]) **(Suppl. Fig. S2, Suppl. Table S2)**. According to estimates based on domain-specific single-copy genes, the obtained MAGs were on average 65.4% complete (minimum: 50.0%, maximum: 100.0%) and 2.7% redundant (minimum: 0.0%, maximum: 9.9%) **(Suppl. Table S2)**. Phylogenomic analyses based on 122 archaeal and 120 bacterial single-copy genes placed the MAGs across 35 bacterial and archaeal phyla according to the GTDB classification [58, 59]. The most represented phyla were Acidobacteriota (n = 172), Actinobacteriota (n = 163), Proteobacteria (Alphaproteobacteria, n = 54; Gammaproteobacteria, n = 39), Chloroflexota (n = 84), and Verrucomicrobiota (n = 43) **(Suppl. Fig. S2)**. Most MAGs (n = 703) belonged to genera that do not comprise formally described species, including 303 MAGs that were placed outside genus-level lineages currently described in GTDB and thus likely represent novel genera **(Suppl. Table S2)**.

To investigate their distribution across the different soil ecosystems, MAGs were dereplicated based on a 99% ANI threshold, yielding a set of 761 non-redundant MAGs **(Fig. 2)**. On average, 15.8% of the reads from each sample were recruited by the set of non-redundant MAGs (minimum: 7.6%, maximum: 30.5%). In agreement with the read-based assessment, we observed differences in MAG composition across the soil ecosystems, with only 50 MAGs shared between the heathland, meadow, and fen soils **(Suppl. Fig. S3a)**. Fen soils harboured the highest number of MAGs, with an average of 155 MAGs per sample **(Suppl. Fig. S3b)**. Although barren and fen soils had similar taxonomic richness according to the read-based estimates, only a small number of MAGs was detected in the barren soils (average of four MAGs per sample). This is likely a result of limited sampling and sequencing of this ecosystem, which consisted of four samples and a total of 7.9 Gb of metagenomic data **(Suppl. Table S1)**. The number of MAGs in heathland and meadow soils was similar (average of 47 and 63 MAGs per sample, respectively) **(Suppl. Fig. S3b)**. In general, barren, heathland, and meadow soils were dominated by the same set of MAGs **(Suppl. Fig. S3c)**. These included members of the Acidobacteriota (*Sulfotelmatobacter* and unclassified genera in the class Acidobacteriae), Actinobacteriota (*Mycobacterium* and unclassified genera in the family Streptosporangiaceae), and Proteobacteria (Alphaproteobacteria: *Reyranella*, *Bradyrhizobium*, and unclassified Xanthobacteraceae; Gammaproteobacteria: unclassified Steroidobacteraceae). On the other hand, fen soils were dominated by MAGs that were not assigned to formally described genera, including lineages of Acidobacteriota (family Koribacteraceae), Actinobacteriota (family Solirubrobacteraceae), Chloroflexota (class Ellin6529), Desulfobacterota (order Desulfobaccales), and Halobacterota.

**Fig. 2.**
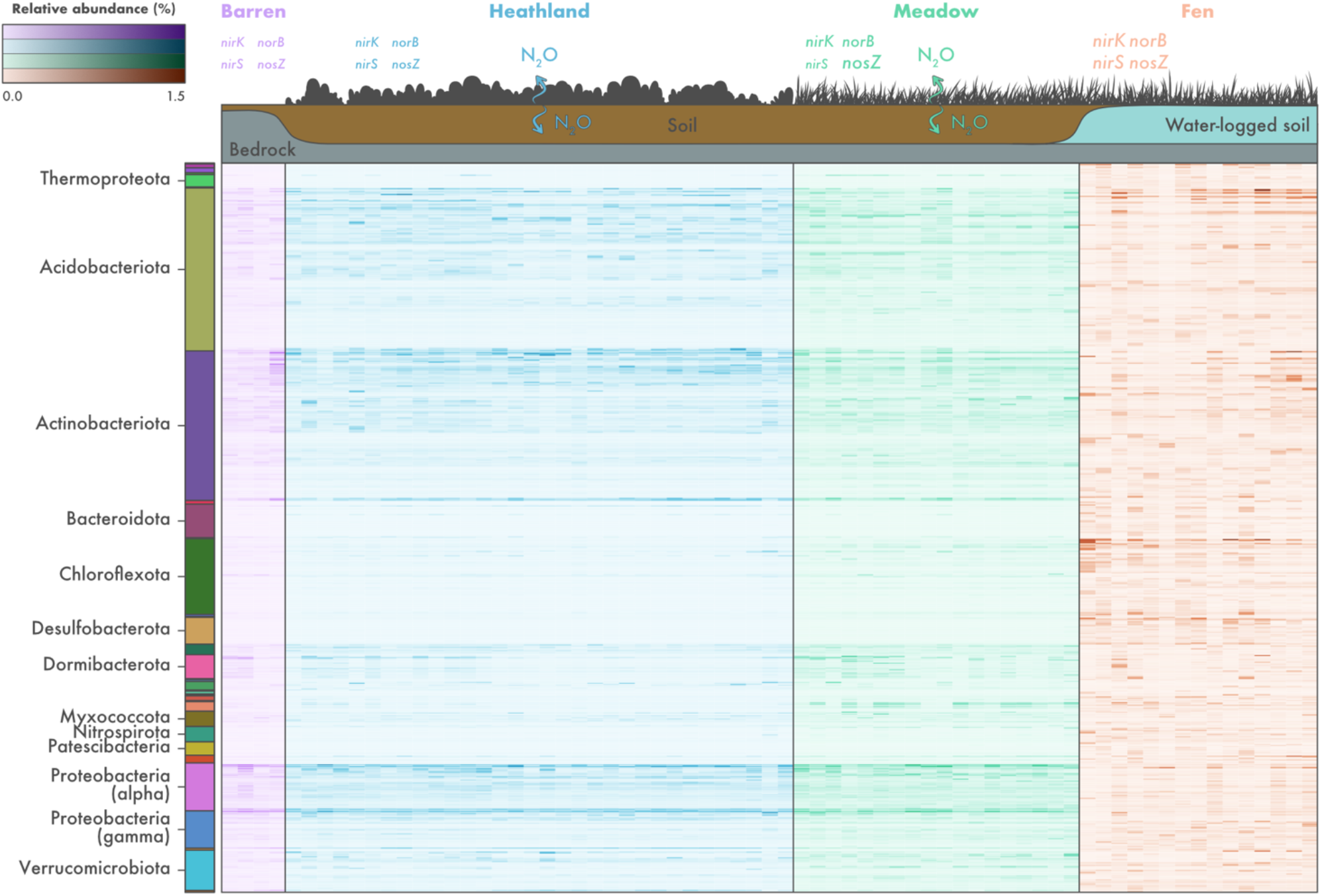
Microbial community composition across different soil ecosystems in the tundra. Relative abundance of 761 non-redundant metagenome-assembled genomes (MAGs) recovered from soils in Kilpisjärvi, northern Finland. Relative abundances were computed as a proportion of the reads mapping to each MAG. Phylum-level taxonomic assignments are shown for the major groups found. The scheme on the top of the figure represents ecosystem-level nitrous oxide (N_2_O) fluxes based on *in situ* measurements **(Fig. 1)** and the abundance of denitrification genes based on a gene-centric analysis **(Suppl. Fig. S1)**. The font size of denitrification genes represents their abundance across the different ecosystems.

### Microorganisms from tundra soils have truncated denitrification pathways

To gain insights into the microorganisms involved with the cycling of N_2_O in tundra soils, we traced the curated denitrification genes to the set of recovered MAGs. Denitrification genes were found in 110 of the 796 MAGs (13.8%) **(Suppl. Table S2)**. These were affiliated with the archaeal phylum Thermoproteota and many bacterial phyla such as Proteobacteria (classes Gamma- and Alphaproteobacteria), Acidobacteriota, Bacteroidota, Actinobacteriota, Chloroflexota, and Verrucomicrobiota **(Fig. 3a)**. However, only 17 MAGs were assigned to a validly described genera **(Suppl. Table S2)**. These included members of the Acidobacteriota (*Solibacter*, *Sulfotelmatobacter*, *Terracidiphilus*, and *Gaiella*), Myxococcota (*Anaeromyxobacter*), Planctomycetota (*Singulisphaera*), Proteobacteria (Alphaproteobacteria: *Bauldia*, *Bradyrhizobium*, *Methylocella*, and *Reyranella*; Gammaproteobacteria: *Gallionella* and *Rhizobacter*), and Verrucomicrobiota (*Lacunisphaera* and *Opitutus*). On average, 1.8% of the reads in each sample were recruited by all denitrifiers combined (minimum: 0.4%, maximum: 6.1%). In general, denitrifiers were most abundant in the fens (1.0–6.1%) and least abundant in the heathlands (0.4–2.1%).

**Fig. 3.**
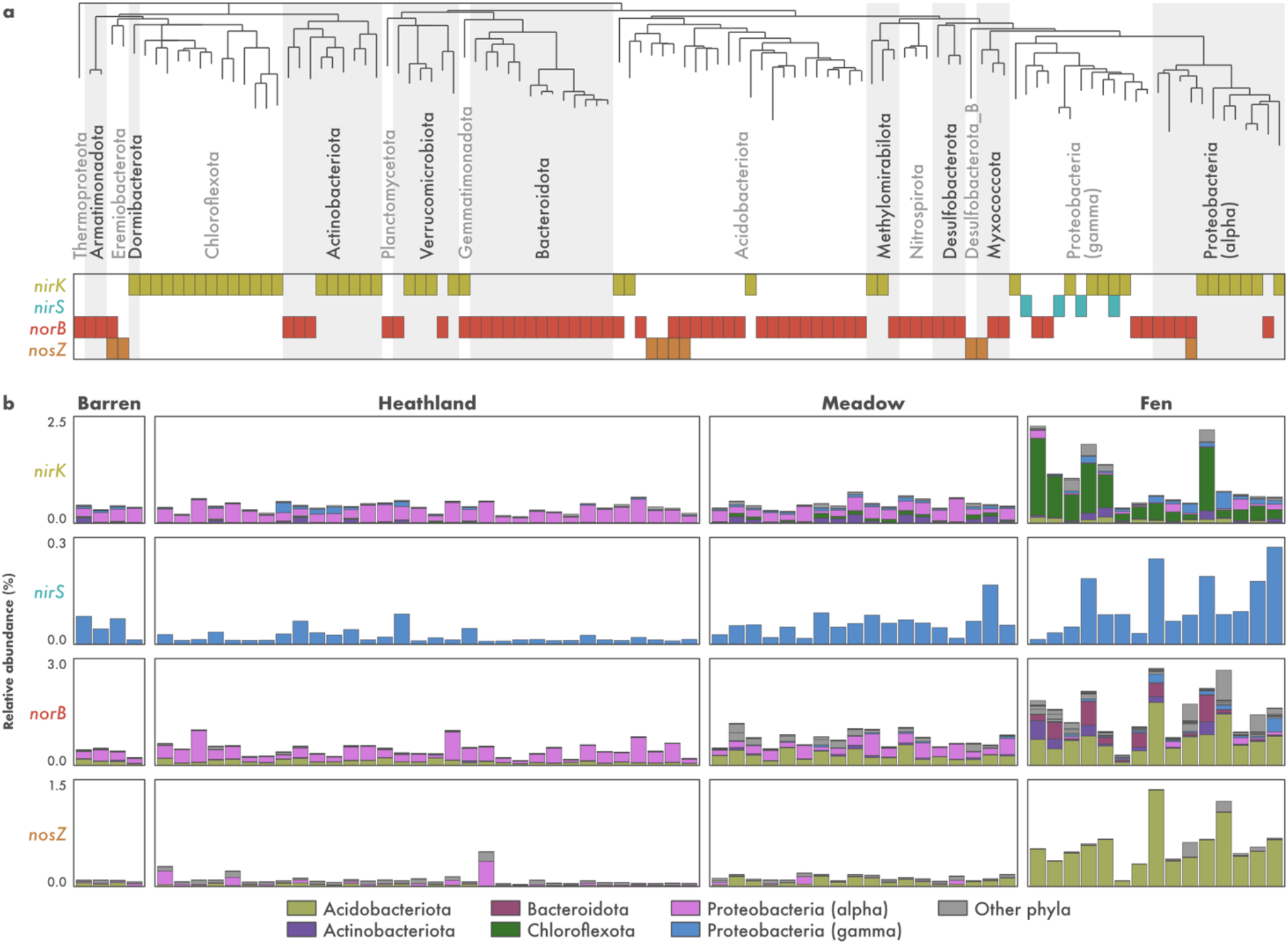
Metabolic potential for denitrification in tundra soils. **a)** Distribution of denitrification genes across 110 metagenome-assembled genomes (MAGs) recovered from tundra soils in Kilpisjärvi, northern Finland. Genes encoding the nitrite (*nirK/nirS*), nitric oxide (*norB*), and nitrous oxide (*nosZ*) reductases were annotated using a three-step approach including (1) identification using hidden Markov models from the KOfam database, (2) manual inspection for the presence of conserved residues at positions associated with the binding of co-factors and active sites, and (3) phylogenetic analyses along with sequences from archaeal and bacterial genomes **(Suppl. Fig. S5)**. **b)** Phylum-level relative abundance of microorganisms harbouring denitrification genes across the different soil ecosystems, computed as a proportion of reads mapping to each MAG.

Genes involved in denitrification were found exclusively in MAGs with truncated denitrification pathways, i.e., MAGs missing one or more genes involved in the complete denitrification process **(Fig. 3a)**. Of the 110 MAGs harbouring denitrification genes, the vast majority (n = 104) encoded only one of the Nir, Nor, and Nos enzymes and no MAG encoded all the three enzymes required for the reduction of NO_2_- to N_2_. Unsurprisingly, co-occurrence of genes encoding the three enzymes was also not observed in any of the other genomic bins of lower quality that were discarded from the final MAG dataset (i.e., bins that were < 50% complete and/or > 10% redundant). To verify if microorganisms with truncated denitrification pathways are common in other tundra systems, we expanded our analysis to 1529 MAGs recovered from permafrost peatland, bog, and fen soils in Stordalen Mire, northern Sweden [37]. Among these, 225 MAGs (14.7%) contained denitrification genes **(Suppl. Fig. S4)**. MAGs encompassed a similar taxonomic profile as observed in the Kilpisjärvi dataset, and MAGs with truncated denitrification pathways were also the norm in Stordalen Mire soils. Only one MAG, assigned to the Gammaproteobacteria genus *Janthinobacterium*, encoded all the Nir, Nor, and Nos enzymes required for the reduction of NO_2_^−^ to N_2_.

### Microorganisms affiliated with the Chloroflexota lineage Ellin6529 are the main denitrifiers *stricto sensu* in fen soils

The reduction of NO_2_- to NO, performed by microorganisms harbouring the *nirK* or *nirS* genes, is the hallmark step of denitrification and is often referred to as denitrification *stricto sensu* as it involves the conversion of a soluble substrate to a gaseous product thus leading to the removal of N from the system [12]. Of the 110 Kilpisjärvi MAGs harbouring genes involved in denitrification, 46 contained *nirK*/*nirS* genes and are thus potential denitrifiers *stricto sensu* **(Fig. 3a)**. These belonged mainly to the bacterial phyla Chloroflexota, Actinobacteriota, and Proteobacteria (classes Alpha- and Gammaproteobacteria). Most MAGs (n = 43) contained the *nirK* gene, which encodes the copper-containing form of Nir **(Suppl. Fig. S5a)**. The *nirS* gene encoding the cytochrome cd1-containing form of Nir was present in four Gammaproteobacteria MAGs **(Suppl. Fig. S5b)**, including one MAG that contained both genes.

The composition of potential denitrifier *stricto sensu* communities differed across the ecosystems **(Fig. 3b)**. MAGs belonging to the Alphaproteobacteria class of the Proteobacteria were the most abundant in the barren, heathland, and meadow soils, particularly the MAG KUL-0154 assigned to the genera *Bradyrhizobium* **(Fig. 4)**. Two other Alphaproteobacteria MAGs that do not correspond to formally described genera in the families Acetobacteraceae and Beijerinckiaceae (KUL-0057 and KUL-0056, respectively) were also found at high abundances. In addition, one Actinobacteriota MAG assigned to an uncharacterized genus in the family Gaiellaceae (KWL-0073), was abundant in the meadow soils. On the other hand, fen communities were dominated by MAGs belonging to the phylum Chloroflexota **(Fig. 3b)**, which included seven MAGs assigned to the class-level lineage Ellin6529 **(Fig. 4)**.

**Fig. 4.**
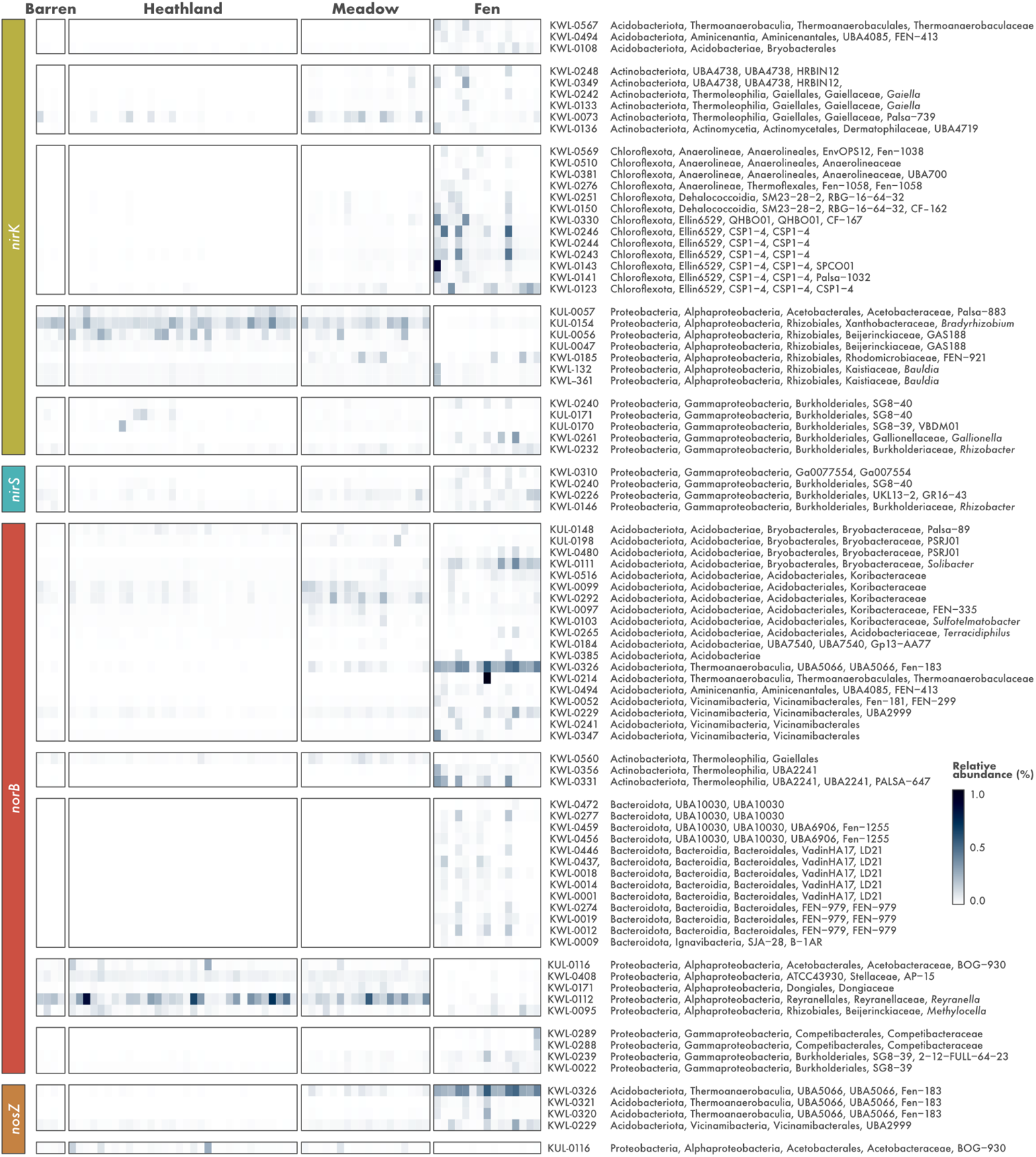
Relative abundance of metagenome-assembled genomes (MAGs) harbouring denitrification genes across different soil ecosystems in the tundra. MAGs were recovered from soils in Kilpisjärvi, northern Finland, and annotated for genes encoding the nitrite (*nirK/nirS*), nitric oxide (*norB*), and nitrous oxide (*nosZ*) reductases using a three-step approach. Relative abundances were computed as a proportion of reads mapping to each MAG. MAG taxonomy is based on the Genome Taxonomy Database (GTDB) release 05-RS95.

None of the Ellin6529 MAGs that were dominant in the fen communities contained the key genes involved in autotrophic carbon fixation, dissimilatory sulfate reduction, dissimilatory nitrate reduction to ammonia, and nitrogen fixation **(Suppl. Table S3)**. Analysis of genes encoding terminal oxidases involved in the aerobic respiratory electron chain revealed that all seven Ellin6529 MAGs harboured the *coxABC* genes encoding the *aa3*-type cytochrome c oxidase. Four MAGs also contained the *cydAB* genes encoding the cytochrome *bd* ubiquinol oxidase, a terminal oxidase with high affinity for oxygen that also plays a role in preventing the inactivation of oxygen-sensitive enzymes and protecting against oxidative and nitrosative stress, toxic compounds such as cyanide, and other stress conditions such as high temperature and high pH [79, 80]. The dominant MAGs in the barren, heathland, and meadow soils encoded a different set of aerobic terminal oxidases. In addition to the *cydAB* genes, the MAGs KUL-0057 and KUL-0154 also contained the *cyoABCD* genes encoding the cytochrome o ubiquinol oxidase, which is the main terminal oxidase under highly aerobic conditions [81], and KUL-0057 also contained genes encoding the *cbb3-type* cytochrome c oxidase, a terminal oxidase with high affinity for oxygen [82]. Genes involved in the Calvin cycle (e.g., *rbcL, rbcS*, and *prkB*) were found in the *Bradyrhizobium* MAG (KUL-0154), and none of the key genes for autotrophic carbon fixation pathways were present in the other Alphaproteobacteria MAGs that were dominant in the barren, heathland, and meadow soils.

### Acidobacteriota with the potential to reduce NO and N_2_O are abundant in the fens

The stepwise reduction of NO to N_2_O and N_2_ carried out by microorganisms containing the *norB* and *nosZ* genes, respectively, represents the final step of denitrification and the main biotic control on N_2_O emissions. Soil denitrification rates depend on multiple environmental conditions such as adequate moisture and inorganic N availability, but whether it results in the emission of N_2_O or N_2_ is ultimately linked to a balance between the activity of NO and N_2_O reducers [11, 15]. *norB* and *nosZ* genes were identified in 62 and 9 Kilpisjärvi MAGs, respectively, belonging mostly to the phyla Actinobacteriota, Bacteroidota, Acidobacteriota, and Proteobacteria (class Alphaproteobacteria) **(Fig. 3a)**. Apart from one Gemmatimonadota and one Acidobacteriota MAG, *norB*- and *nosZ*-containing MAGs were almost exclusively non-denitrifiers *stricto sensu*, i.e., they did not harbour the *nirK/nirS* genes involved in the reduction of NO_2_- to NO. Most MAGs (n = 48) harboured a *norB* gene encoding the monomeric, quinol-dependent form of Nor (qNor), while the remaining MAGs (n = 8) encoded the cytochrome c-dependent Nor (cNor) **(Suppl. Fig. S5c)**. In regards to the *nosZ* gene, most MAGs (n = 6) contained sequences affiliated with the clade II (also known as atypical) NosZ [14, 15, 27] **(Suppl. Fig. S5d)**. Only four MAGs contained both the *norB* and *nosZ* genes and thus have the potential to reduce NO completely to N_2_ **(Fig. 3a)**.

As observed for the denitrifier *stricto sensu* communities, the communities of potential NO and N_2_O reducers also differed between the ecosystems **(Fig. 3b)**. MAGs assigned to the Alphaproteobacteria class of the Proteobacteria were the most abundant in the barren, heathland, and meadow soils. In particular, the MAG KWL-0112 assigned to the genera *Reyranella* was the dominant *norB*-containing MAG, while KUL-0116 (belonging to an uncharacterized genus in the family Acetobacteraceae) was the dominant MAG harbouring the *nosZ* gene **(Fig. 4)**. On the other hand, fen communities were dominated by Acidobacteriota MAGs **(Fig. 3b)**, particularly the *norB*- and *nosZ*-containing MAG KWL-0326 affiliated with the class Thermoanaerobaculia **(Fig. 4)**. This MAG contained the same set of genes encoding aerobic terminal oxidases as found in the *nirK*-containing Ellin6529 MAGs that were dominant in the fen sites, namely *coxABC* and *cydAB* **(Suppl. Table S3)**. No genes involved in carbon fixation, dissimilatory sulfate reduction, dissimilatory nitrate reduction to ammonia, and nitrogen fixation were found in any of the dominant *norB*- and *nosZ*-containing MAGs.

To elucidate the phylogenetic placement of the Acidobacteriota MAGs and to verify if the potential for NO and N_2_O reduction is present in other members of this phylum, we analysed all available genomes of Acidobacteriota strains and candidate taxa available on GenBank. This revealed that genes encoding the Nir and Nos enzymes are widespread across the phylum Acidobacteriota **(Fig. 5)**. Genes encoding the Nor enzyme were present in all but one of the six Acidobacteriota subdivisions with genomes from cultured representatives. This included the strains *Acidobacterium ailaaui* PMMR2 (subdivision Gp1), *Acidipila* sp. 4G-K13 (Gp1), *Silvibacterium bohemicum* DSM 103733 and *S*. *bohemicum* S15 (Gp1), Acidobacteriaceae bacterium URHE0068 (Gp1), *Edaphobacter aggregans* DSM 19364 (Gp1), *Luteitalea pratensis* DSM 100886 (Gp6), *Geothrix fermentans* DSM 14018 (Gp8), and *Thermoanaerobaculum aquaticum* MP-01 (Gp23), as well as the candidate taxa *Candidatus* Koribacter versatilis Ellin345 (Gp1), *Candidatus* Sulfotelmatomonas gaucii SbA5 (Gp1), and *Candidatus* Solibacter usitatus Ellin6076 (Gp3). On the other hand, genes encoding the Nos enzyme were found only in members of the subdivisions Gp6 and Gp23.

**Fig. 5.**
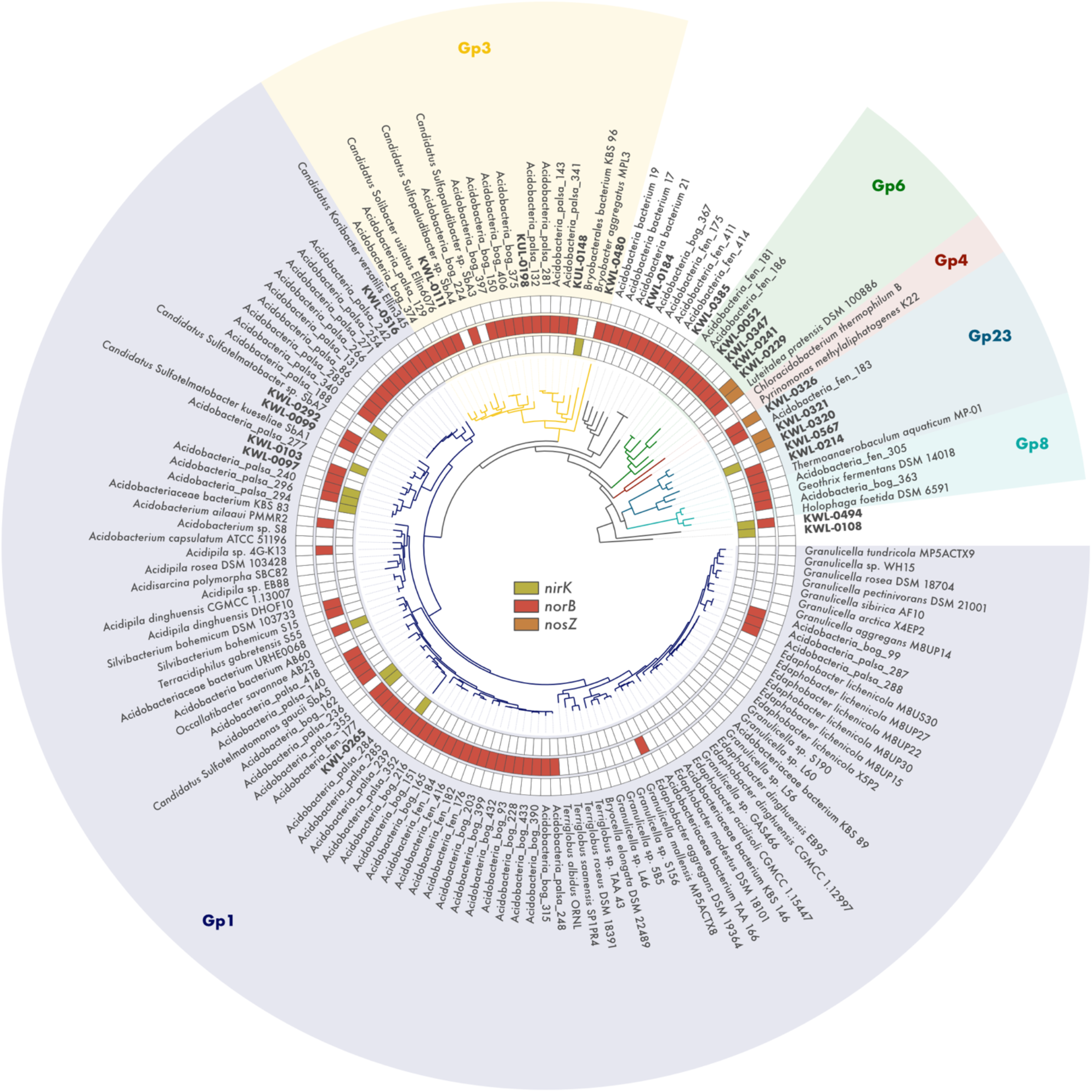
Metabolic potential for denitrification among members of the phylum Acidobacteriota. Phylogenomic analysis of 85 Acidobacteriota metagenome-assembled genomes (MAGs) containing denitrification genes recovered from tundra soils in Kilpisjärvi (northern Finland) and Stordalen Mire (northern Sweden), and 69 genomes of Acidobacteriota strains and candidate taxa. Maximum likelihood tree based on concatenated alignments of 23 ribosomal proteins and rooted with *Escherichia coli* ATCC 11775 (not shown). Genes encoding the nitrite (*nirK*), nitric oxide (*norB*), and nitrous oxide (*nosZ*) reductases were annotated using a three-step approach to avoid false positives (see methods).

## Discussion

The 796 MAGs obtained in the present study by a manual binning and curation effort represent one of the largest genomic catalogues of microorganisms from tundra soils to date. Earlier gene-centric investigations have revealed the potential for complete denitrification in tundra soils [22, 83], however, these approaches fail to reveal the wider genomic context of the genes involved in this pathway. By applying the genome-resolved metagenomics approach, we traced denitrification genes to specific microbial populations, thereby allowing a detailed investigation of the genomic makeup of potential denitrifiers in tundra soils. This approach also enabled us to access the genomes of uncultured, poorly characterized taxa, which comprise the majority of the microorganisms in soils and other complex ecosystems [33, 34].

Our genome-resolved survey revealed that denitrification across different tundra soil ecosystems is dominated by microorganisms with truncated denitrification pathways (i.e., harbouring only a subset of the genes required for complete denitrification), most of which represent poorly characterized taxa without cultured representatives. The congruence of these findings in both our original dataset of northern Finland soils and a re-analysis of a comprehensive metagenomic dataset from soils in northern Sweden [37] suggests that truncated denitrification pathways are not a methodological artifact arising from the metabolic reconstruction of fragmented genomes. Indeed, recent genome-resolved investigations have shown that cross-feeding between microorganisms with truncated metabolic pathways, also known as metabolic handoffs, are the norm across a wide range of ecosystems such as grassland soil, aquifer sediment, groundwater, and the ocean, and not only in relation to denitrification but other redox transformations as well [30, 84, 85]. Although it has been established that denitrification is a community effort performed by different microbial populations [12–15], these genome-resolved metagenomic studies are beginning to reveal a more in-depth, ecosystem-centric representation of the denitrification pathway. In addition to their predominance in genomic databases [14], it appears that truncated denitrifiers are also dominant within defined ecosystems across various terrestrial and aquatic biomes, including the tundra. It has been suggested that the partitioning of metabolic pathways across different populations via metabolic handoffs is advantageous as it eliminates competition between enzymes accelerating substrate consumption [15, 31] and provides flexibility and resilience to the communities in face of environmental disturbances [30]. We further hypothesize that the predominance of denitrification pathways characterized mostly by metabolic handoffs in tundra soils could be related to N limitation. If metabolic handoffs enable a more effective substrate consumption as previously suggested [15, 31], truncated denitrification pathways would be favoured in tundra soils which are mostly N limited but undergo rapid surges in N availability, e.g., during the spring melting season [86].

Tundra ecosystems are typically heterogeneous. Previous studies in the Kilpisjärvi region have shown that soil properties such as pH and moisture do not have any strong relationship with the macrotopography of the area (50–500 m scale). Instead, environmental variation is controlled by the fine-scale mesotopographic variation of the relief (2–20 m scale), resulting in a mosaic of different soil ecosystems with contrasting vegetation [40, 75–78]. Our results showed that denitrifier communities in the tundra differ between drier upland ecosystems (barren, heathland, and meadow soils) and water-logged fens. This is likely related to differences in soil moisture affecting oxygen availability in these ecosystems. The dominant denitrifier populations in the oxic dry upland soils, related to the genera *Bradyrhizobium*, *Reyranella*, and other uncharacterized genera in the class Alphaproteobacteria, encoded aerobic terminal oxidases that are active under highly aerobic conditions as well as oxidases with high oxygen affinity [81, 82]. The former likely provides an adaptive advantage in these soils by allowing rapid aerobic growth under standard conditions of high oxygen availability, and the latter would sustain growth in microoxic niches within the soil matrix and during periods of reduced oxygen availability (e.g., during the spring melting season).

On the other hand, fen soils are continuously inundated because they are located at lower topographic positions where the water table is permanently at or near the soil surface. The result is a mostly anoxic environment due to the slow rate at which oxygen diffuses into the water-logged soil, favouring reduced rather than oxidized soil chemistry. In line with this, we found a predominance of anaerobic processes in the fens, including a higher abundance of genes involved in denitrification, sulfate reduction, and methanogenesis, the latter supported by *in situ* measurements showing net CH_4_ emission at the fen sites. Communities of potential denitrifiers in the fen soils were dominated by somewhat enigmatic taxa, namely potential NO_2_- reducers affiliated with the class Ellin6529 of the Chloroflexota and NO/N_2_O reducers assigned to the subdivision Gp23 of the Acidobacteriota. Both groups are major members of microbial communities in soils worldwide [87], and RNA-based investigations have shown that they are active in tundra soils during both summer and winter seasons [40, 88]. *Thermoanaerobaculum aquaticum* MP-01, the only cultivated member of the Acidobacteriota subdivision Gp23, is a strictly anaerobic bacterium that has been shown to use Fe and Mn, but not NO_3_^-^ nor NO_2_-, as electron acceptors in anaerobic respiration [89]. However, studies investigating the use of nitrogen oxides in anaerobic respiration usually provide soluble NO_3_^-^ or NO_2_- as electron acceptors, not the gases NO and N_2_O, which bias against truncated denitrifiers that do not contain the *narG* and *nirK*/*nirS* genes [90]. Ellin6529 – formerly G04 – were first detected by culture-independent methods in alpine tundra wet meadow soil in the Colorado Rocky Mountains, USA [91], and later isolated in a study targeting slow-growing and mini-colony forming bacteria from Australian agricultural soil [92]. However, their ecological, physiological, and metabolic preferences remain largely unknown. Their genomic composition and high abundance in the water-logged, anoxic fen soils suggest that the Ellin6529 and Gp23 populations found in this study are likely able to grow anaerobically with the use of NO and N_2_O as electron acceptors. However, it is known that in addition to their role in anaerobic respiration, NO and N_2_O reduction can be used as a detoxification mechanism or as electron sink for metabolism. For example, the aerobe *Gemmatimonas aurantica* T-27 is not able to grow on N_2_O alone, but can use N_2_O as electron acceptor transiently when oxygen is depleted [93].

In addition to microbial community structure, differences in N_2_O fluxes observed between upland and fen soils also appear to be linked to soil moisture. Some of the drier upland sites investigated were hotspots of N_2_O consumption. This is particularly interesting for the acidic heathland soils, as low pH is known to impair the expression of the NosZ enzyme thus promoting N_2_O emission [94, 95]. On the other hand, fens had close to net-zero N_2_O fluxes, which is in line with previous observations for water-saturated soils both in the tundra [7] and worldwide [11, 13]. This has been linked to lower rates of N mineralization and nitrification in anoxic ecosystems, which limit the availability of NO_3_^-^ and NO_2_- and promote complete denitrification, resulting in N_2_ as end product rather than N_2_O. Indeed, supplementing fen soils in the tundra with NO_3_^-^ and NO_2_- has shown to promote N_2_O emissions [96]. Moreover, climate change models predict lowering of the water table in high-latitude wetlands, which could lead to increased N_2_O emissions from these ecosystems which contain substantial amounts of both C and N bound to the soil organic matter [97, 98].

## Conclusions

A better understanding of denitrification is paramount for our ability to model N_2_O emissions and mitigate climate change. High-latitude environments in particular have experienced amplified warming in recent decades, a trend that is likely to continue in the coming centuries. As mechanisms of GHG emissions are very climate sensitive, the contribution of tundra soils to global GHG atmospheric levels is thus predicted to increase in the future leading to a positive feedback loop. Compared with CO_2_ and CH_4_, measurements of N_2_O fluxes in tundra soils are sparse and are rarely coupled with a characterization of the microorganisms involved, making the magnitude and drivers of N_2_O fluxes across the polar regions uncertain. While microorganisms with truncated denitrification pathways appear to dominate the denitrifier communities investigated here, the potential for complete denitrification was present at the ecosystem level. In addition to a better monitoring of N_2_O emissions throughout the tundra biome, our results suggest that a better understanding of the contribution of tundra soil to global N_2_O levels relies on the elucidation of the regulatory mechanisms of metabolic handoffs in communities dominated by truncated denitrifiers.

## Declarations

### Ethics approval and consent to participate

Not applicable.

### Consent for publication

Not applicable.

### Availability of data and materials

Raw metagenomic data and assembled MAGs have been submitted to the European Nucleotide Archive (ENA) under the project PRJEB41762. All the code used can be found in https://github.com/ArcticMicrobialEcology/Kilpisjarvi-MAGs.

### Competing interests

The authors declare that they have no competing interests.

### Funding

This work was funded by the Academy of Finland (grants 314114 and 335354) and the University of Helsinki. SV was funded by the Microbiology and Biotechnology Doctoral Programme (MBDP). AMV was funded by the Academy of Finland (grant 286950), the Otto Malm Foundation, and the Gordon and Betty Moore Foundation (grant 8414). MEM was supported by the Academy of Finland (grants 314630 and 317054).

### Author contributions

JH and ML designed the research; SV and JH performed nucleic acid extraction and metagenomic library preparation; AMV and MEM designed and performed the GHG flux measurements and analyses; ISP analysed the data and wrote the manuscript; EER and TOD contributed with the analyses; all authors contributed to the final version of the manuscript.

## Acknowledgements

We would like to acknowledge CSC – IT Centre for Science for providing the necessary computing resources and Kimmo Mattila for IT support; the staff from the Kilpisjärvi Biological Station, Tanja Orpana, Aino Rutanen, Anniina Sarekoski, Johanna Kerttula, and the members of the BioGeoClimate Modelling Lab for assistance with fieldwork and soil characterization; Jillian Banfield and Christina Biasi for helpful discussion; Laura Cappelatti for proof-reading the manuscript; Murat Eren, Sebastian Lücker, Donovan Parks, and Antonios Kioukis for tips, recommendations, and troubleshooting; and all anonymous reviewers who provided important insights to the original manuscript.

## Supplementary Figures

**Suppl. Fig. S1.**
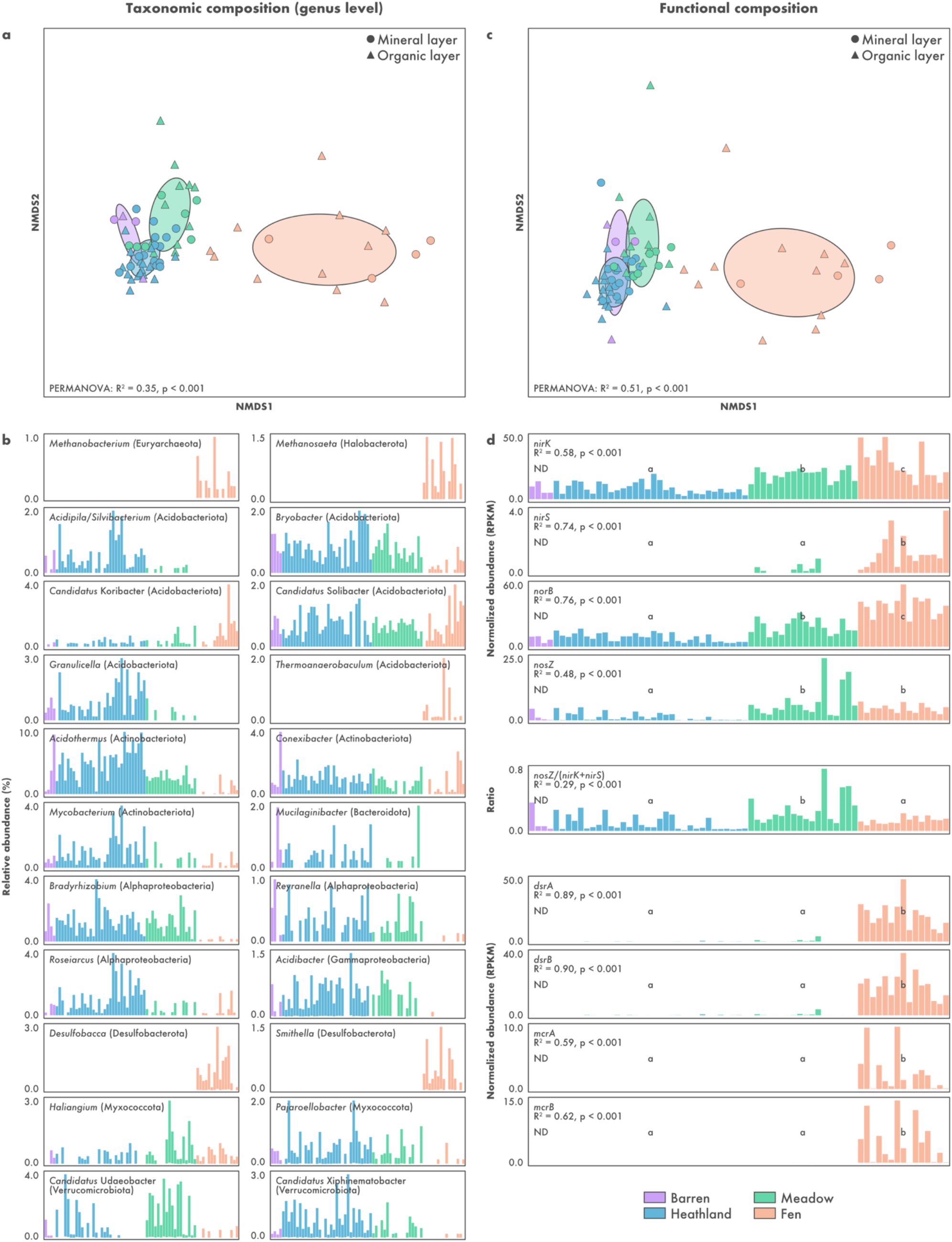
The microbial diversity of Kilpisjärvi soils as seen using a gene-centric approach. Taxonomic composition was computed based on the annotation of unassembled SSU rRNA gene sequences against the SILVA database. Functional annotation was done by searching assembled genes against the KOfam database. The annotation of putative denitrification genes was confirmed using a phylogenetic approach. **a, c)** Non-metric multidimensional scaling (NMDS) of taxonomic and functional community structure, respectively. Differences between the ecosystems were assessed using permutational ANOVA (PERMANOVA). **b)** Abundance profile of the five most abundant genera in each ecosystem. **d)** Abundance profile of marker genes for denitrification, sulfate reduction, and methanogenesis. Ecosystems followed by different letters are significantly different (one-way ANOVA, p < 0.05). Samples from barren soils were not included in the ANOVA procedure due to the limited number of samples (ND: not determined).

**Suppl. Fig. S2.**
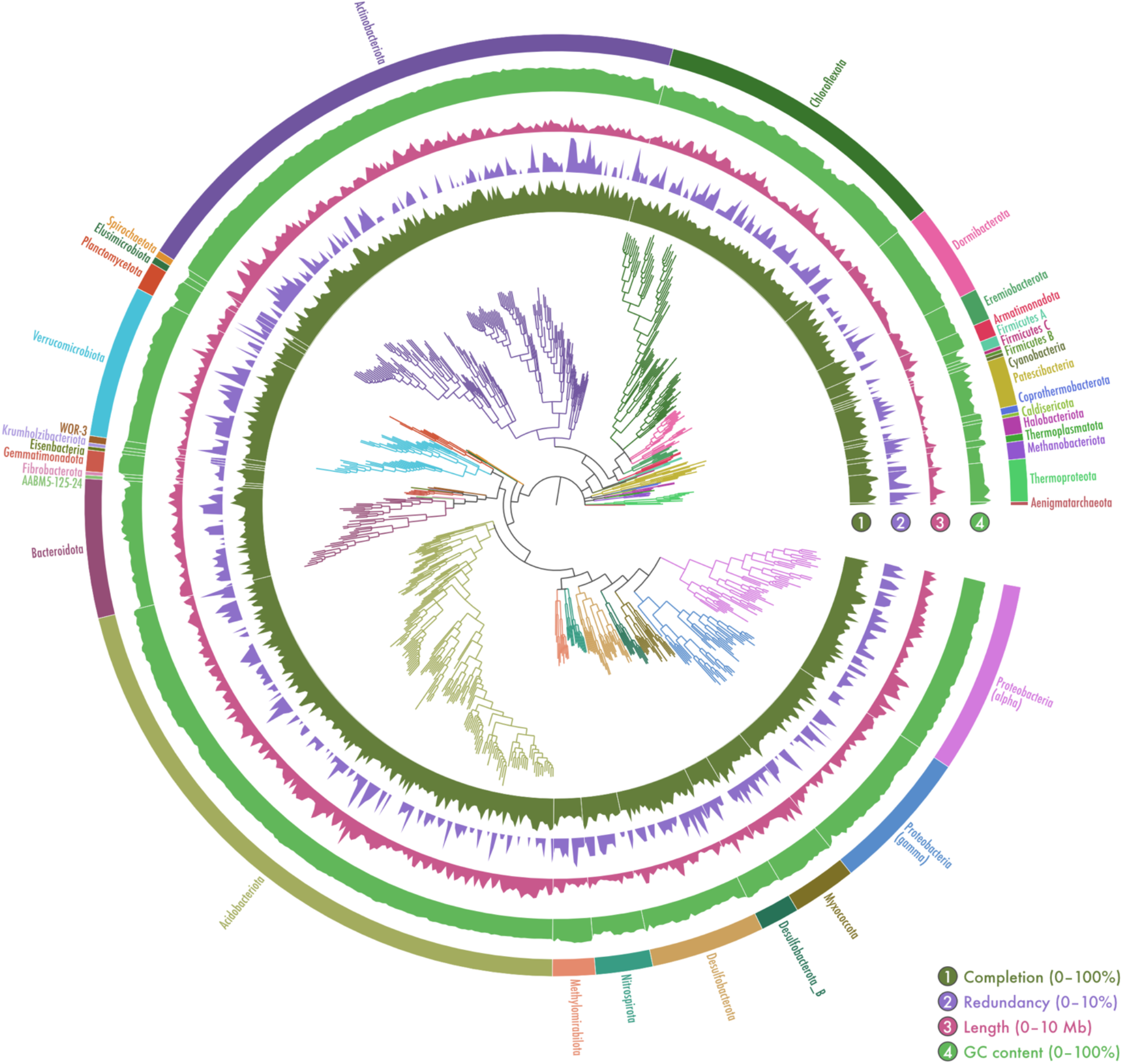
Genome-resolved metagenomics of tundra soils. Phylogenomic placement and assembly statistics of 796 metagenome-assembled genomes (MAGs) recovered from soils in Kilpisjärvi, northern Finland. Unrooted maximum likelihood tree based on concatenated alignments of amino acid sequences from 122 archaeal and 120 bacterial single-copy genes. Additional information about the MAGs can be found in **Suppl. Table S2**.

**Suppl. Fig. S3.**
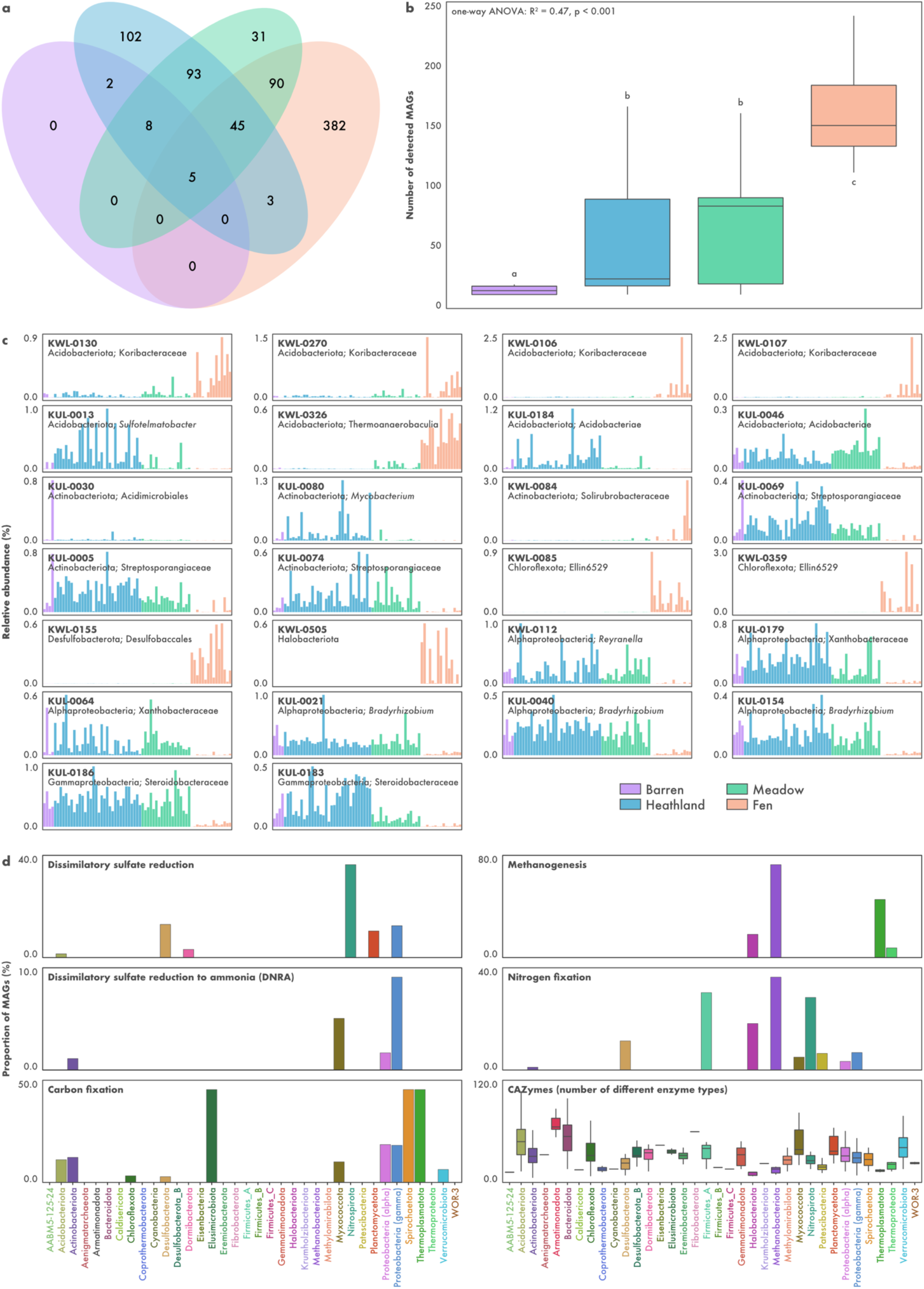
Overview of the microbial diversity in Kilpisjärvi soils based on a genome-resolved approach. a) Number of metagenome-assembled genomes (MAGs) shared between the different ecosystems. **b)** Number of detected MAGs across the ecosystems. **c)** Relative abundance of the ten most abundant MAGs in each ecosystem, computed as a proportion of reads mapping to each MAG. **d)** Metabolic potential of the MAGs based on the annotation of genes against the KOfam database. Bar plots represent the proportion of MAGs in each phylum with complete pathways, i.e., containing ≥ 75% of the genes in the pathway. Boxplots of carbohydrate-active enzymes (CAZymes) show the number of different enzyme types identified in each MAG.

**Suppl. Fig. S4.**
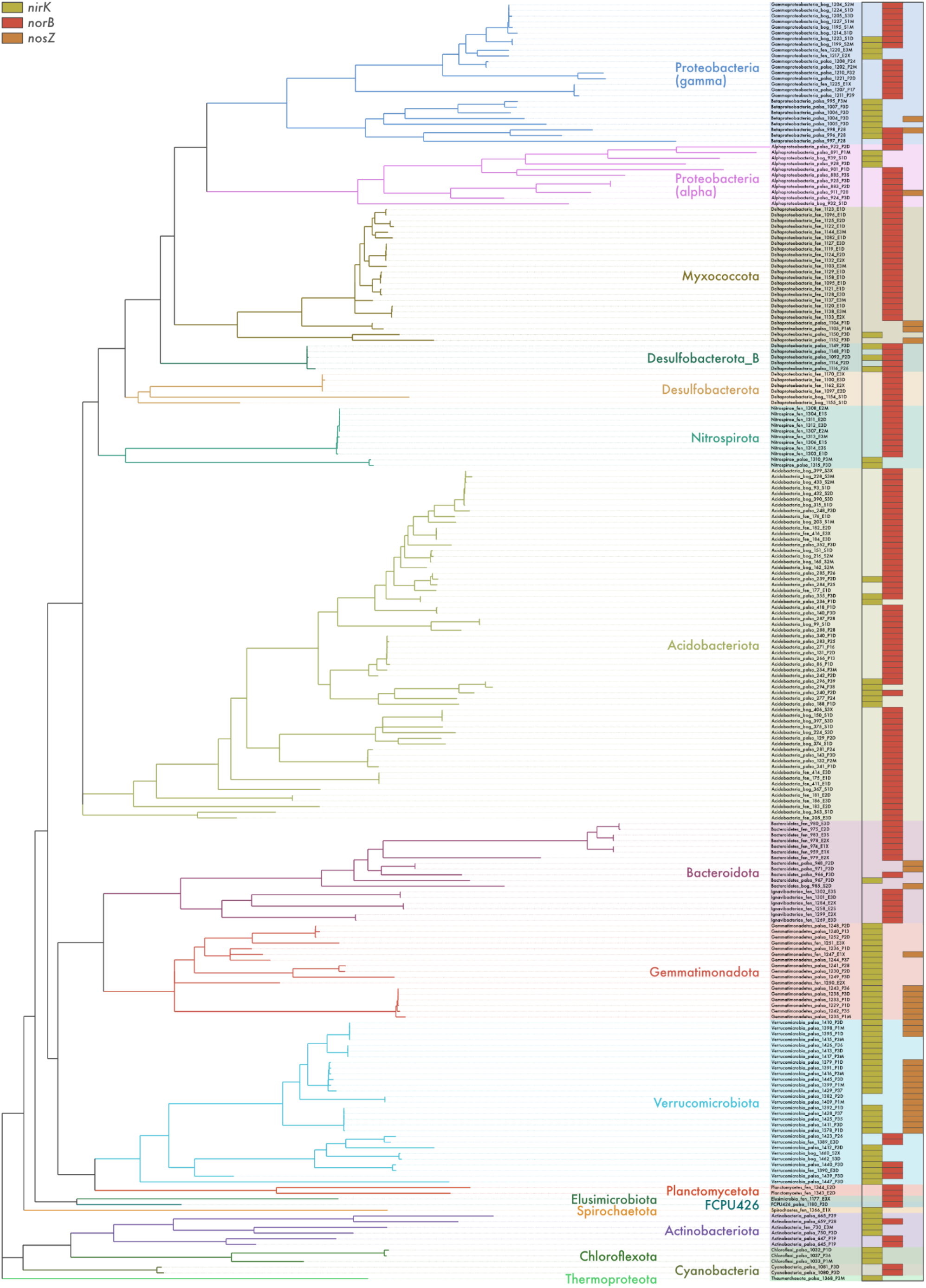
Metabolic potential for denitrification in Stordalen Mire soils. Distribution of denitrification genes across 225 metagenome-assembled genomes (MAGs) from permafrost peatland, bog, and fen soils in Stordalen Mire, northern Sweden. Genes encoding the nitrite (*nirK*), nitric oxide (*norB*), and nitrous oxide (*nosZ*) reductases were annotated using a three-step approach including (1) identification using hidden Markov models from the KOfam database, (2) manual inspection for the presence of conserved residues at positions associated with the binding of co-factors and active sites, and (3) phylogenetic analyses along with sequences from archaeal and bacterial genomes. Phylogenomic analysis of MAGs was done based on concatenated alignments of amino acid sequences from 122 archaeal and 120 bacterial single-copy genes.

**Suppl. Fig. S5.**
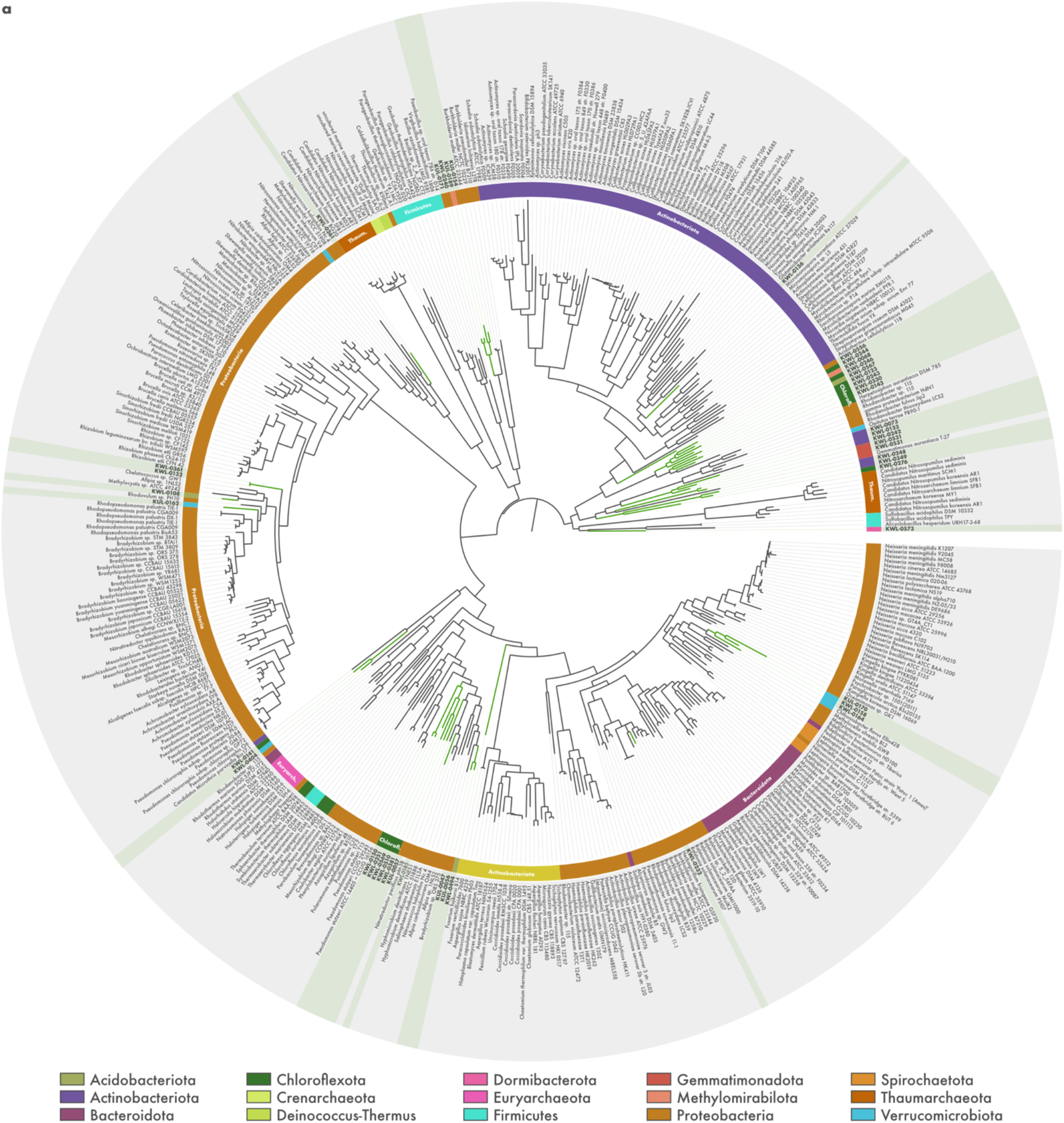
Phylogeny of a) *nirK*, b) *nirS*, c) *norB*, and d) *nosZ* sequences from metagenome-assembled genomes (MAGs) recovered from tundra soils in Kilpisjärvi, northern Finland. Midpoint-rooted maximum-likelihood trees of translated sequences from Kilpisjärvi MAGs (highlighted) along with reference sequences from archaeal and bacterial genomes.

**Suppl. Fig. S5.**
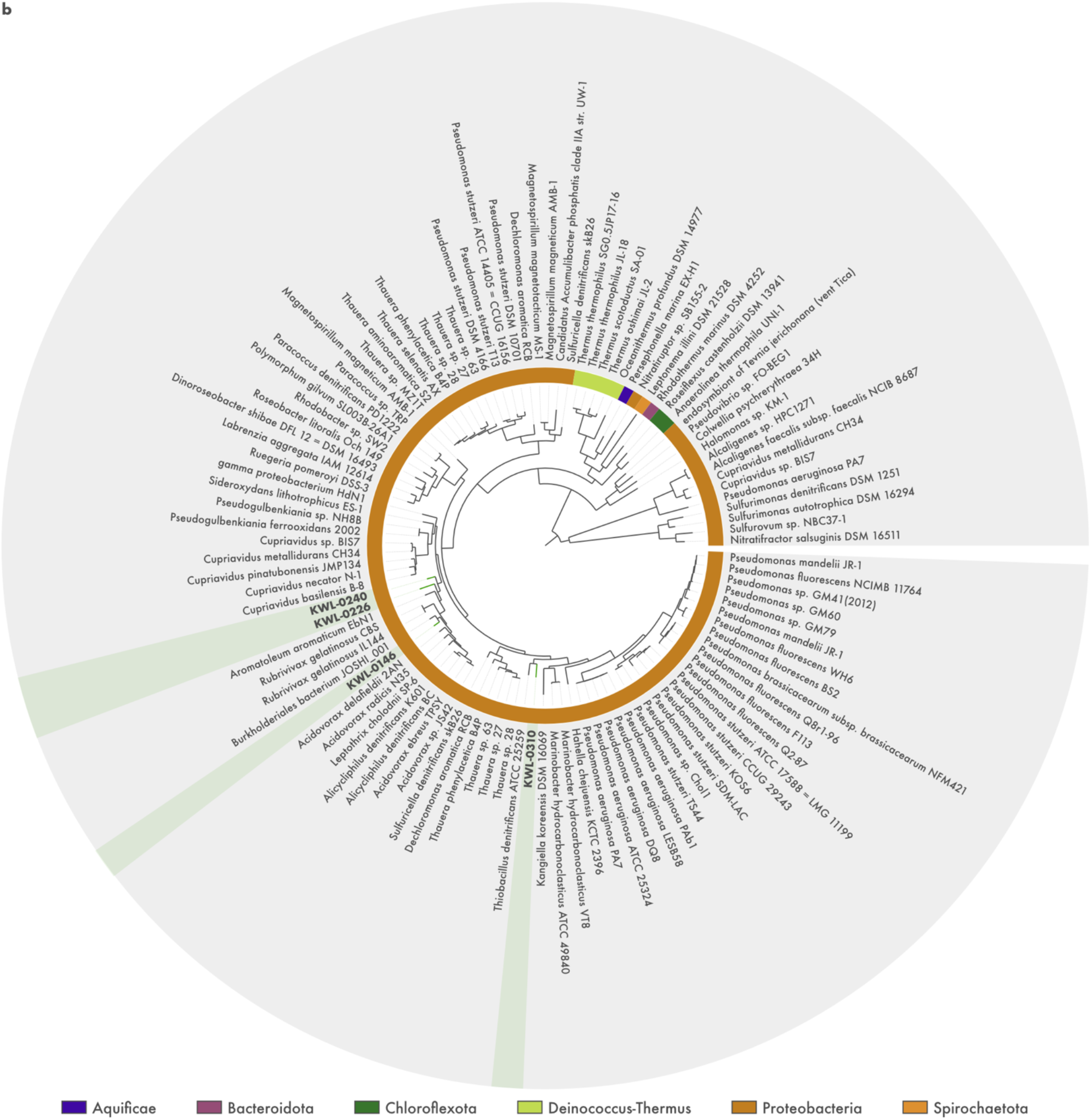
Phylogeny of a) *nirK*, b) *nirS*, c) *norB*, and d) *nosZ* sequences from metagenome-assembled genomes (MAGs) recovered from tundra soils in Kilpisjärvi, northern Finland. Midpoint-rooted maximum-likelihood trees of translated sequences from Kilpisjärvi MAGs (highlighted) along with reference sequences from archaeal and bacterial genomes.

**Suppl. Fig. S5.**
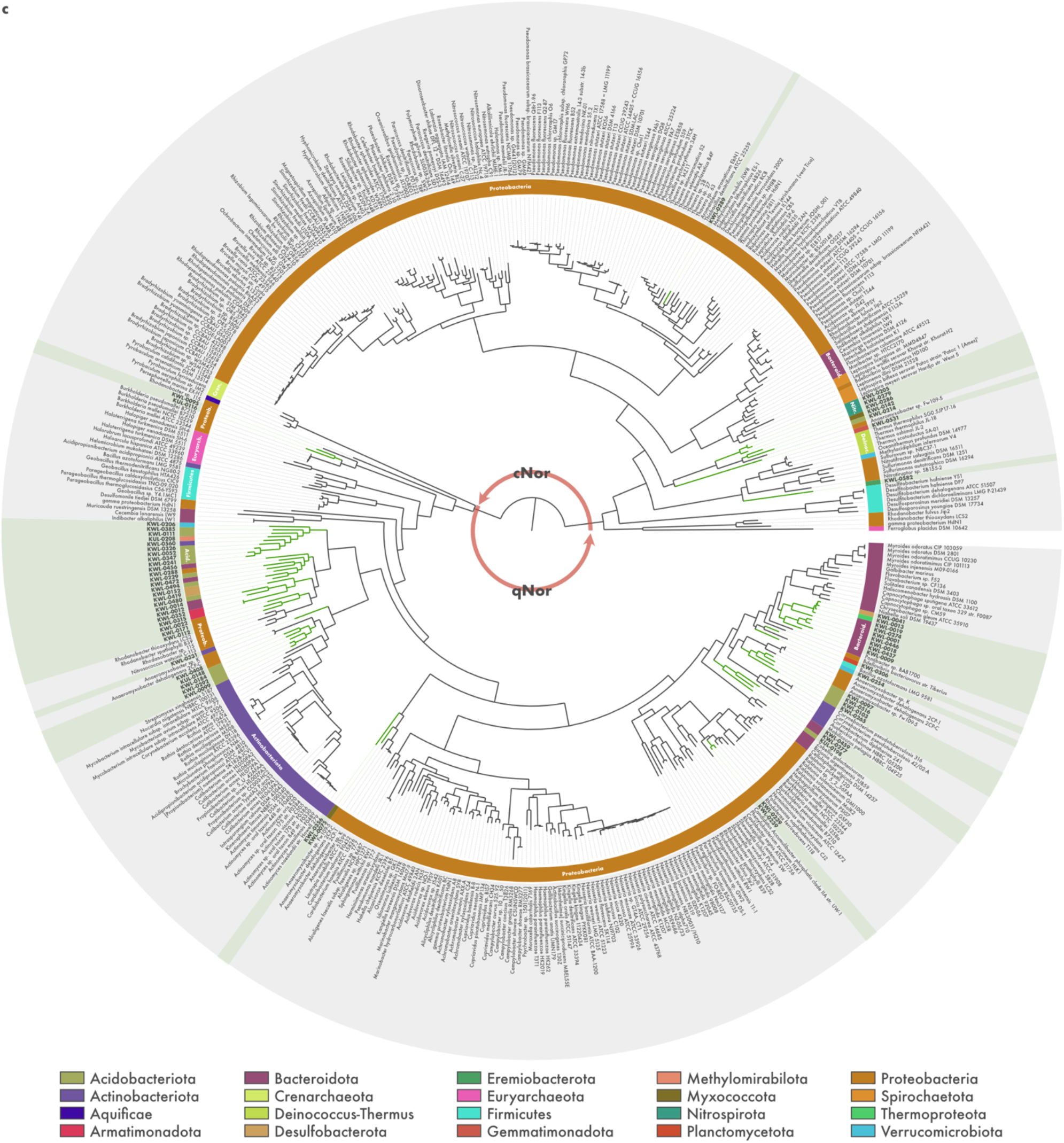
Phylogeny of a) *nirK*, b) *nirS*, c) *norB*, and d) *nosZ* sequences from metagenome-assembled genomes (MAGs) recovered from tundra soils in Kilpisjärvi, northern Finland. Midpoint-rooted maximum-likelihood trees of translated sequences from Kilpisjärvi MAGs (highlighted) along with reference sequences from archaeal and bacterial genomes.

**Suppl. Fig. S5.**
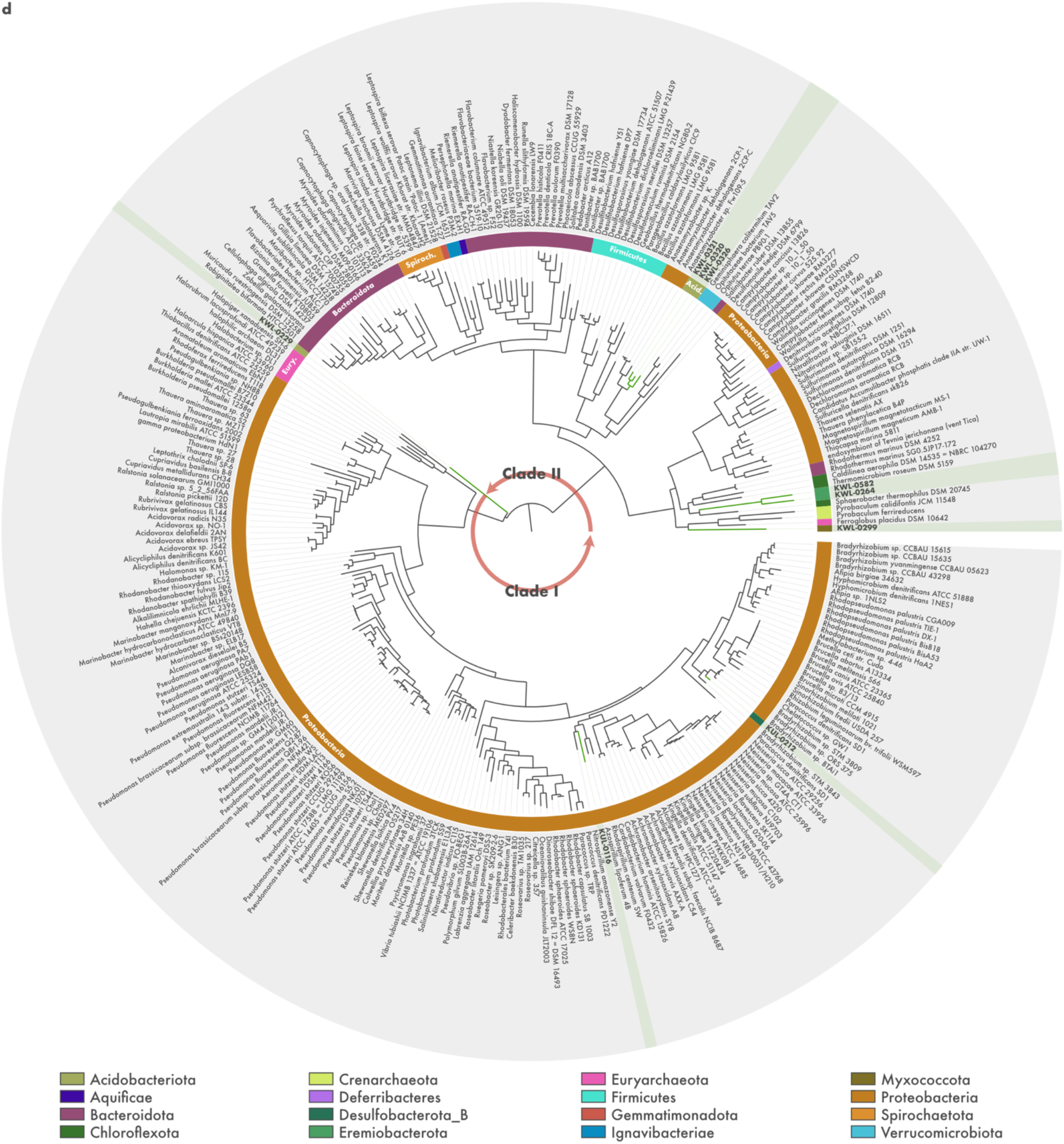
Phylogeny of a) *nirK*, b) *nirS*, c) *norB*, and d) *nosZ* sequences from metagenome-assembled genomes (MAGs) recovered from tundra soils in Kilpisjärvi, northern Finland. Midpoint-rooted maximum-likelihood trees of translated sequences from Kilpisjärvi MAGs (highlighted) along with reference sequences from archaeal and bacterial genomes.

## Notes

### Competing Interest Statement

The authors have declared no competing interest.

## References

1. IPCC, editor. Climate Change 2013: The Physical Science Basis. Contribution of Working Group I to the Fifth Assessment Report of the Intergovernmental Panel on Climate Change. Cambridge: Cambridge University Press; 2013.

2. Tian H, Xu R, Canadell JG, Thompson RL, Winiwarter W, Suntharalingam P, et al. A comprehensive quantification of global nitrous oxide sources and sinks. Nature. 2020;586:248–56.

3. Repo ME, Susiluoto S, Lind SE, Jokinen S, Elsakov V, Biasi C, et al. Large N2O emissions from cryoturbated peat soil in tundra. Nat Geosci. 2009;2:189–92.

4. Marushchak ME, Pitkämäki A, Koponen H, Biasi C, Seppälä M, Martikainen PJ. Hot spots for nitrous oxide emissions found in different types of permafrost peatlands. Glob Change Biol. 2011;17:2601–14.

5. Stewart KJ, Grogan P, Coxson DS, Siciliano SD. Topography as a key factor driving atmospheric nitrogen exchanges in arctic terrestrial ecosystems. Soil Biol Biochem. 2014;70:96–112.

6. Voigt C, Marushchak ME, Lamprecht RE, Jackowicz-Korczyński M, Lindgren A, Mastepanov M, et al. Increased nitrous oxide emissions from Arctic peatlands after permafrost thaw. Proc Natl Acad Sci. 2017;114:6238–43.

7. Voigt C, Marushchak ME, Abbott BW, Biasi C, Elberling B, Siciliano SD, et al. Nitrous oxide emissions from permafrost-affected soils. Nat Rev Earth Environ. 2020;1:420–34.

8. Schuur EAG, McGuire AD, Schädel C, Grosse G, Harden JW, Hayes DJ, et al. Climate change and the permafrost carbon feedback. Nature. 2015;520:171–9.

9. Hugelius G, Loisel J, Chadburn S, Jackson RB, Jones M, MacDonald G, et al. Large stocks of peatland carbon and nitrogen are vulnerable to permafrost thaw. Proc Natl Acad Sci. 2020;117:20438–46.

10. Post E, Alley RB, Christensen TR, Macias-Fauria M, Forbes BC, Gooseff MN, et al. The polar regions in a 2°C warmer world. Sci Adv. 2019;5:eaaw9883.

11. Butterbach-Bahl K, Baggs EM, Dannenmann M, Kiese R, Zechmeister-Boltenstern S. Nitrous oxide emissions from soils: how well do we understand the processes and their controls? Philos Trans R Soc B Biol Sci. 2013;368:20130122.

12. Zumft WG. Cell biology and molecular basis of denitrification. Microbiol Mol Biol Rev MMBR. 1997;61:533–616.

13. Wallenstein MD, Myrold DD, Firestone M, Voytek M. Environmental controls on denitrifying communities and denitrification rates: insights from molecular methods. Ecol Appl. 2006;16:2143–52.

14. Graf DRH, Jones CM, Hallin S. Intergenomic Comparisons Highlight Modularity of the Denitrification Pathway and Underpin the Importance of Community Structure for N2O Emissions. PLoS ONE. 2014;9:e114118.

15. Hallin S, Philippot L, Löffler FE, Sanford RA, Jones CM. Genomics and Ecology of Novel N2O-Reducing Microorganisms. Trends Microbiol. 2018;26:43–55.

16. Liu X-Y, Koba K, Koyama LA, Hobbie SE, Weiss MS, Inagaki Y, et al. Nitrate is an important nitrogen source for Arctic tundra plants. Proc Natl Acad Sci. 2018;115:3398–403.

17. Kou D, Yang G, Li F, Feng X, Zhang D, Mao C, et al. Progressive nitrogen limitation across the Tibetan alpine permafrost region. Nat Commun. 2020;11:3331.

18. Yergeau E, Kang S, He Z, Zhou J, Kowalchuk GA. Functional microarray analysis of nitrogen and carbon cycling genes across an Antarctic latitudinal transect. ISME J. 2007;1:163–79.

19. Yergeau E, Hogues H, Whyte LG, Greer CW. The functional potential of high Arctic permafrost revealed by metagenomic sequencing, qPCR and microarray analyses. ISME J. 2010;4:1206–14.

20. Palmer K, Biasi C, Horn MA. Contrasting denitrifier communities relate to contrasting N2O emission patterns from acidic peat soils in arctic tundra. ISME J. 2012;6:1058–77.

21. Dai H-T, Zhu R-B, Sun B-W, Che C-S, Hou L-J. Effects of Sea Animal Activities on Tundra Soil Denitrification and nirS-and nirK-Encoding Denitrifier Community in Maritime Antarctica. Front Microbiol. 2020;11:573302.

22. Ortiz M, Bosch J, Coclet C, Johnson J, Lebre P, Salawu-Rotimi A, et al. Microbial Nitrogen Cycling in Antarctic Soils. Microorganisms. 2020;8:1442.

23. Brummell ME, Farrell RE, Siciliano SD. Greenhouse gas soil production and surface fluxes at a high arctic polar oasis. Soil Biol Biochem. 2012;52:1–12.

24. Chapuis-Lardy L, Wrage N, Metay A, Chotte J-L, Bernoux M. Soils, a sink for N_2_ O? A review. Glob Change Biol. 2007;13:1–17.

25. Bakken LR, Bergaust L, Liu B, Frostegård Å. Regulation of denitrification at the cellular level: a clue to the understanding of N_2_O emissions from soils. Philos Trans R Soc B Biol Sci. 2012;367:1226–34.

26. Philippot L, Andert J, Jones CM, Bru D, Hallin S. Importance of denitrifiers lacking the genes encoding the nitrous oxide reductase for N2O emissions from soil: role of denitrifier diversity for N2O fluxes. Glob Change Biol. 2011;17:1497–504.

27. Sanford RA, Wagner DD, Wu Q, Chee-Sanford JC, Thomas SH, Cruz-Garcia C, et al. Unexpected nondenitrifier nitrous oxide reductase gene diversity and abundance in soils. Proc Natl Acad Sci. 2012;109:19709–14.

28. Jones CM, Graf DR, Bru D, Philippot L, Hallin S. The unaccounted yet abundant nitrous oxide-reducing microbial community: a potential nitrous oxide sink. ISME J. 2013;7:417–26.

29. Jones CM, Spor A, Brennan FP, Breuil M-C, Bru D, Lemanceau P, et al. Recently identified microbial guild mediates soil N2O sink capacity. Nat Clim Change. 2014;4:801–5.

30. Anantharaman K, Brown CT, Hug LA, Sharon I, Castelle CJ, Probst AJ, et al. Thousands of microbial genomes shed light on interconnected biogeochemical processes in an aquifer system. Nat Commun. 2016;7:13219.

31. Lilja EE, Johnson DR. Segregating metabolic processes into different microbial cells accelerates the consumption of inhibitory substrates. ISME J. 2016;10:1568–78.

32. Yu T, Zhuang Q. Quantifying global N2O emissions from natural ecosystem soils using trait-based biogeochemistry models. Biogeosciences. 2019;16:207–22.

33. Rappé MS, Giovannoni SJ. The Uncultured Microbial Majority. Annu Rev Microbiol. 2003;57:369–94.

34. Steen AD, Crits-Christoph A, Carini P, DeAngelis KM, Fierer N, Lloyd KG, et al. High proportions of bacteria and archaea across most biomes remain uncultured. ISME J. 2019;13:3126–30.

35. Mackelprang R, Waldrop MP, DeAngelis KM, David MM, Chavarria KL, Blazewicz SJ, et al. Metagenomic analysis of a permafrost microbial community reveals a rapid response to thaw. Nature. 2011;480:368–71.

36. Hultman J, Waldrop MP, Mackelprang R, David MM, McFarland J, Blazewicz SJ, et al. Multi-omics of permafrost, active layer and thermokarst bog soil microbiomes. Nature. 2015;521:208–12.

37. Woodcroft BJ, Singleton CM, Boyd JA, Evans PN, Emerson JB, Zayed AAF, et al. Genome-centric view of carbon processing in thawing permafrost. Nature. 2018;560:49–54.

38. Pirinen P, Simola H, Aalto J, Kaukoranta J-P, Karlsson P, Ruuhela R. Climatological statistics of Finland 1981-2010. Helsinki: Finnish Meteorological Institute; 2012.

39. Livingston GP, Hutchinson GL. Enclosure-based measurement of trace gas exchange: applications and sources of error. In: Biogenic trace gases: Measuring emissions from soil and water. Oxford, United Kingdom: Blackwell Science. p. 14–51.

40. Viitamäki S, Pessi IS, Virkkala A-M, Niittynen P, Kemppinen J, Eronen-Rasimus E, et al. The activity and functions of subarctic soil microbial communities vary across vegetation types. bioRxiv. 2021. https://doi.org/10.1101/2021.06.12.448001.

41. Ewels P, Magnusson M, Lundin S, Käller M. MultiQC: summarize analysis results for multiple tools and samples in a single report. Bioinformatics. 2016;32:3047–8.

42. Martin M. Cutadapt removes adapter sequences from high-throughput sequencing reads. EMBnet.journal. 2011;17:10.

43. Leger A, Leonardi T. pycoQC, interactive quality control for Oxford Nanopore Sequencing. J Open Source Softw. 2019;4:1236.

44. Bengtsson-Palme J, Hartmann M, Eriksson KM, Pal C, Thorell K, Larsson DGJ, et al. METAXA 2: improved identification and taxonomic classification of small and large subunit rRNA in metagenomic data. Mol Ecol Resour. 2015;15:1403–14.

45. Quast C, Pruesse E, Yilmaz P, Gerken J, Schweer T, Yarza P, et al. The SILVA ribosomal RNA gene database project: improved data processing and web-based tools. Nucleic Acids Res. 2012;41:D590–6.

46. Schloss PD, Westcott SL, Ryabin T, Hall JR, Hartmann M, Hollister EB, et al. Introducing mothur: Open-Source, Platform-Independent, Community-Supported Software for Describing and Comparing Microbial Communities. Appl Environ Microbiol. 2009;75:7537–41.

47. Wang Q, Garrity GM, Tiedje JM, Cole JR. Naïve Bayesian Classifier for Rapid Assignment of rRNA Sequences into the New Bacterial Taxonomy. Appl Environ Microbiol. 2007;73:5261–7.

48. Li D, Liu C-M, Luo R, Sadakane K, Lam T-W. MEGAHIT: an ultra-fast single-node solution for large and complex metagenomics assembly via succinct de Bruijn graph. Bioinformatics. 2015;31:1674–6.

49. Kolmogorov M, Bickhart DM, Behsaz B, Gurevich A, Rayko M, Shin SB, et al. metaFlye: scalable long-read metagenome assembly using repeat graphs. Nat Methods. 2020. https://doi.org/10.1038/s41592-020-00971-x.

50. Langmead B, Salzberg SL. Fast gapped-read alignment with Bowtie 2. Nat Methods. 2012;9:357–9.

51. Li H, Handsaker B, Wysoker A, Fennell T, Ruan J, Homer N, et al. The Sequence Alignment/Map format and SAMtools. Bioinformatics. 2009;25:2078–9.

52. Walker BJ, Abeel T, Shea T, Priest M, Abouelliel A, Sakthikumar S, et al. Pilon: An Integrated Tool for Comprehensive Microbial Variant Detection and Genome Assembly Improvement. PLoS ONE. 2014;9:e112963.

53. Mikheenko A, Saveliev V, Gurevich A. MetaQUAST: evaluation of metagenome assemblies. Bioinformatics. 2016;32:1088–90.

54. Eren AM, Esen ÖC, Quince C, Vineis JH, Morrison HG, Sogin ML, et al. Anvi’o: an advanced analysis and visualization platform for ‘omics data. PeerJ. 2015;3:e1319.

55. Hyatt D, Chen G-L, LoCascio PF, Land ML, Larimer FW, Hauser LJ. Prodigal: prokaryotic gene recognition and translation initiation site identification. BMC Bioinformatics. 2010;11:119.

56. Eddy SR. Accelerated Profile HMM Searches. PLoS Comput Biol. 2011;7:e1002195.

57. Buchfink B, Xie C, Huson DH. Fast and sensitive protein alignment using DIAMOND. Nat Methods. 2015;12:59–60.

58. Parks DH, Chuvochina M, Waite DW, Rinke C, Skarshewski A, Chaumeil P-A, et al. A standardized bacterial taxonomy based on genome phylogeny substantially revises the tree of life. Nat Biotechnol. 2018;36:996–1004.

59. Parks DH, Chuvochina M, Chaumeil P-A, Rinke C, Mussig AJ, Hugenholtz P. A complete domain-to-species taxonomy for Bacteria and Archaea. Nat Biotechnol. 2020. https://doi.org/10.1038/s41587-020-0501-8.

60. Alneberg J, Bjarnason BS, de Bruijn I, Schirmer M, Quick J, Ijaz UZ, et al. Binning metagenomic contigs by coverage and composition. Nat Methods. 2014;11:1144–6.

61. Bowers RM, Kyrpides NC, Stepanauskas R, Harmon-Smith M, Doud D, Reddy TBK, et al. Minimum information about a single amplified genome (MISAG) and a metagenome-assembled genome (MIMAG) of bacteria and archaea. Nat Biotechnol. 2017;35:725–31.

62. Aramaki T, Blanc-Mathieu R, Endo H, Ohkubo K, Kanehisa M, Goto S, et al. KofamKOALA: KEGG Ortholog assignment based on profile HMM and adaptive score threshold. Bioinformatics. 2020;36:2251–2.

63. Katoh K, Standley DM. MAFFT Multiple Sequence Alignment Software Version 7: Improvements in Performance and Usability. Mol Biol Evol. 2013;30:772–80.

64. Okonechnikov K, Golosova O, Fursov M. Unipro UGENE: a unified bioinformatics toolkit. Bioinformatics. 2012;28:1166–7.

65. Decleyre H, Heylen K, Tytgat B, Willems A. Highly diverse nirK genes comprise two major clades that harbour ammonium-producing denitrifiers. BMC Genomics. 2016;17:155.

66. Li Y, Bali S, Borg S, Katzmann E, Ferguson SJ, Schuler D. Cytochrome cd1 Nitrite Reductase NirS Is Involved in Anaerobic Magnetite Biomineralization in Magnetospirillum gryphiswaldense and Requires NirN for Proper d1 Heme Assembly. J Bacteriol. 2013;195:4297–309.

67. Heylen K, Keltjens J. Redundancy and modularity in membrane-associated dissimilatory nitrate reduction in Bacillus. Front Microbiol. 2012;3.

68. Price MN, Dehal PS, Arkin AP. FastTree 2 – Approximately Maximum-Likelihood Trees for Large Alignments. PLoS ONE. 2010;5:e9490.

69. Woodcroft BJ. CoverM. 2021.

70. Li H. Minimap and miniasm: fast mapping and de novo assembly for noisy long sequences. Bioinformatics. 2016;32:2103–10.

71. Chaumeil P-A, Mussig AJ, Hugenholtz P, Parks DH. GTDB-Tk: a toolkit to classify genomes with the Genome Taxonomy Database. Bioinformatics. 2019;:btz848.

72. Edgar RC. MUSCLE: multiple sequence alignment with high accuracy and high throughput. Nucleic Acids Res. 2004;32:1792–7.

73. Zhang H, Yohe T, Huang L, Entwistle S, Wu P, Yang Z, et al. dbCAN2: a meta server for automated carbohydrate-active enzyme annotation. Nucleic Acids Res. 2018;46:W95–101.

74. Jain C, Rodriguez-R LM, Phillippy AM, Konstantinidis KT, Aluru S. High throughput ANI analysis of 90K prokaryotic genomes reveals clear species boundaries. Nat Commun. 2018;9:5114.

75. le Roux PC, Aalto J, Luoto M. Soil moisture’s underestimated role in climate change impact modelling in low-energy systems. Glob Change Biol. 2013;19:2965–75.

76. Niittynen P, Heikkinen RK, Aalto J, Guisan A, Kemppinen J, Luoto M. Fine-scale tundra vegetation patterns are strongly related to winter thermal conditions. Nat Clim Change. 2020. https://doi.org/10.1038/s41558-020-00916-4.

77. le Roux PC, Pellissier L, Wisz MS, Luoto M. Incorporating dominant species as proxies for biotic interactions strengthens plant community models. J Ecol. 2014;102:767–75.

78. Kemppinen J, Niittynen P, Aalto J, le Roux PC, Luoto M. Water as a resource, stress and disturbance shaping tundra vegetation. Oikos. 2019;128:811–22.

79. Borisov VB, Gennis RB, Hemp J, Verkhovsky MI. The cytochrome bd respiratory oxygen reductases. Biochim Biophys Acta BBA - Bioenerg. 2011;1807:1398–413.

80. Giuffrè A, Borisov VB, Arese M, Sarti P, Forte E. Cytochrome bd oxidase and bacterial tolerance to oxidative and nitrosative stress. Biochim Biophys Acta BBA - Bioenerg. 2014;1837:1178–87.

81. Dinamarca MA, Ruiz-Manzano A, Rojo F. Inactivation of Cytochrome *o* Ubiquinol Oxidase Relieves Catabolic Repression of the *Pseudomonas putida* GPo1 Alkane Degradation Pathway. J Bacteriol. 2002;184:3785–93.

82. Bueno E, Mesa S, Bedmar EJ, Richardson DJ, Delgado MJ. Bacterial Adaptation of Respiration from Oxic to Microoxic and Anoxic Conditions: Redox Control. Antioxid Redox Signal. 2012;16:819–52.

83. Makhalanyane TP, Van Goethem MW, Cowan DA. Microbial diversity and functional capacity in polar soils. Curr Opin Biotechnol. 2016;38:159–66.

84. Diamond S, Andeer PF, Li Z, Crits-Christoph A, Burstein D, Anantharaman K, et al. Mediterranean grassland soil C–N compound turnover is dependent on rainfall and depth, and is mediated by genomically divergent microorganisms. Nat Microbiol. 2019;4:1356–67.

85. Sun X, Ward BB. Novel metagenome-assembled genomes involved in the nitrogen cycle from a Pacific oxygen minimum zone. ISME Commun. 2021;1:26.

86. Westergaard-Nielsen A, Balstrøm T, Treier UA, Normand S, Elberling B. Estimating meltwater retention and associated nitrate redistribution during snowmelt in an Arctic tundra landscape. Environ Res Lett. 2020;15:034025.

87. Delgado-Baquerizo M, Oliverio AM, Brewer TE, Benavent-González A, Eldridge DJ, Bardgett RD, et al. A global atlas of the dominant bacteria found in soil. Science. 2018;359:320–5.

88. Männistö MK, Kurhela E, Tiirola M, Häggblom MM. Acidobacteria dominate the active bacterial communities of Arctic tundra with widely divergent winter-time snow accumulation and soil temperatures. FEMS Microbiol Ecol. 2013;84:47–59.

89. Losey NA, Stevenson BS, Busse H-J, Damsté JSS, Rijpstra WIC, Rudd S, et al. Thermoanaerobaculum aquaticum gen. nov., sp. nov., the first cultivated member of Acidobacteria subdivision 23, isolated from a hot spring. Int J Syst Evol Microbiol. 2013;63 Pt_11:4149–57.

90. Lycus P, Lovise Bøthun K, Bergaust L, Peele Shapleigh J, Reier Bakken L, Frostegård Å. Phenotypic and genotypic richness of denitrifiers revealed by a novel isolation strategy. ISME J. 2017;11:2219–32.

91. Costello EK, Schmidt SK. Microbial diversity in alpine tundra wet meadow soil: novel Chloroflexi from a cold, water-saturated environment. Environ Microbiol. 2006;8:1471–86.

92. Davis KER, Sangwan P, Janssen PH. Acidobacteria, Rubrobacteridae and Chloroflexi are abundant among very slow-growing and mini-colony-forming soil bacteria. Environ Microbiol. 2011;13:798–805.

93. Park D, Kim H, Yoon S. Nitrous Oxide Reduction by an Obligate Aerobic Bacterium, Gemmatimonas aurantiaca Strain T-27. Appl Environ Microbiol. 2017;83.

94. Liu B, Frostegård Å, Bakken LR. Impaired Reduction of N_2_O to N_2_ in Acid Soils Is Due to a Posttranscriptional Interference with the Expression of nosZ. mBio. 2014;5.

95. Samad MS, Biswas A, Bakken LR, Clough TJ, de Klein CAM, Richards KG, et al. Phylogenetic and functional potential links pH and N2O emissions in pasture soils. Sci Rep. 2016;6:35990.

96. Palmer K, Horn MA. Denitrification Activity of a Remarkably Diverse Fen Denitrifier Community in Finnish Lapland Is N-Oxide Limited. PLOS ONE. 2015;10:e0123123.

97. Smith K. The potential for feedback effects induced by global warming on emissions of nitrous oxide by soils. Glob Change Biol. 1997;3:327–38.

98. Kåresdotter E, Destouni G, Ghajarnia N, Hugelius G, Kalantari Z. Mapping the Vulnerability of Arctic Wetlands to Global Warming. Earths Future. 2021;9.

